# The genetic interaction between HIV and the antibody repertoire

**DOI:** 10.1101/646968

**Authors:** Nicolas Strauli, Emily Kathleen Fryer, Olivia Pham, Mohamed Abdel-Mohsen, Shelley N. Facente, Christopher Pilcher, Pleuni Pennings, Satish Pillai, Ryan D. Hernandez

## Abstract

The interaction between human immunodeficiency virus (HIV) and the antibody repertoire (AbR) during chronic infection can provide important information for HIV vaccine research, yet has not been well-characterized on a systems level. We deeply sequenced the HIV population and the AbR of ten HIV-infected, antiretroviral (ART)-naïve individuals, each with 10-20 longitudinal samples spanning 4-14 years. Our unbiased sequencing approach identified partitions of AbRs showing evidence of interaction with autologous HIV populations. We show that these HIV-associated partitions are enriched for the V gene segments of known HIV broadly neutralizing antibodies (bnAbs), indicating that the HIV-responding component of the AbR can be identified via time-series genetic data. Despite this evidence for larger-scale AbR/HIV interactions at the sub-population level, we found little to no evidence for antagonistic coevolution (i.e. an arms race). This suggests that antagonistic coevolution is either rare, or hard to detect, which has important vaccine design implications.

## Introduction

Since the beginning of the modern pandemic in 1981 (Faria et al., 2014; Gottlieb et al., 1981), human immunodeficiency virus (HIV) has been the source of incredible scientific scrutiny. While there has been great progress in the development of antiretroviral therapy (ART), which can now manage the disease indefinitely (albeit only for those who have access to them), a cure remains elusive (Stein, Storcksdieck genannt Bonsmann, & Streeck, 2016), and little progress has been made toward a preventative vaccine (Trovato, D’Apice, Prisco, & De Berardinis, 2018). Prevention efforts have recently made significant headway by implementing preexposure prophylaxis (PrEP) to high risk individuals. However, in its current form, this strategy has its downsides, such as a reliance on daily self-administration, significant financial burden, and health side effects (Weinstein, Yang, & Cohen, 2017). Thus, cure and vaccine strategies remain the elusive goal for HIV research. Together, the distinct gains in treatment, yet relative lack of gains in HIV prevention, has resulted in a stalemate of sorts, where instead of HIV being triumphantly eradicated by modern science, it has settled into a persistent, yet treatable reality of human life.

The fervent hope for progress is particularly palpable in HIV vaccine research. This fervor is mainly fueled by the fact that effective HIV immunity is entirely possible and well documented, as a broadly neutralizing humoral immune response against HIV occurs naturally in 10-20% of those chronically infected (Binley et al., 2008; Hraber et al., 2014; Sather et al., 2009). It seems that while humans universally cannot clear an HIV infection, some can develop immunity to future infections. Moreover, potent humoral immunity has been associated with spontaneous immunological suppression of HIV following disease acquisition in the absence of ART (Freund et al., 2017). If a vaccine were to be designed that could somehow recapitulate the broad and potent anti-HIV humoral immune responses observed in naturally resistant individuals, then effective protective immunity may be achievable. While a good deal of progress has been made in this avenue of research, it has not yet resulted in an effective vaccine. Among the promising discoveries are broadly neutralizing antibodies (bnAbs), which are monoclonal antibodies (Abs) that can single-handedly neutralize up to 90% of heterologous HIV strains (McCoy & Burton, 2017; X. Wu et al., 2011). To study the development of these bnAbs, researchers have utilized a post-hoc deep-sequencing approach by which a set of primers are developed that will preferentially amplify a subset of the antibody repertoire (AbR)—the population of antibodies in an organism—that is known to contain a bnAb lineage (Bhiman et al., 2015; Liao et al., 2013; X. Wu et al., 2011). The advantage of this approach is that one can cut through the incredible noise and complexity of the AbR to focus on a particular lineage of importance. However, such approaches miss the diversity of Ab lineages interacting with HIV that may have important effects on immunity outcomes. To our knowledge, only 2 studies (Hoehn et al., 2015; Setliff et al., 2018) have deeply sequenced the AbR in an unbiased fashion in the context of HIV infection, but they did not collect paired HIV sequence data to directly study genetic interactions.

The common narrative of bnAb development is that they are in a coevolutionary arms race (Dawkins & Krebs, 1979) with the autologous HIV population (Rantalainen et al., 2018; Zhou, Ton, Morriss, Nguyen, & Fera, 2018). This relationship specifically refers to ‘antagonistic coevolution’, which for simplicity, we will here to refer to as ‘coevolution’. There is good reason to suspect that this coevolutionary hypothesis is true: bnAbs tend to be quite derived relative to their inferred naïve ancestors (Bhiman et al., 2015), they tend to take a long time to develop (years), and there tends to be a time dependence on neutralization capabilities (i.e. HIV-neutralizing Abs are more likely to neutralize autologous virus from the past, and less likely to neutralize contemporaneous or future autologous virus) (Gray et al., 2011; Liao et al., 2013). However, there is also evidence contrary to the arms race hypothesis: bnAbs can arise relatively quickly, and with few mutations (Nicole A. Doria-Rose et al., 2014; Goo, Chohan, Nduati, & Overbaugh, 2014; MacLeod et al., 2016; Simonich et al., 2016), and superinfections—multiple HIV infections in the same individual—don’t necessarily drive further evolution in existing HIV-neutralizing Ab lineages, but rather promote the development of *de novo* HIV-targeting Ab lineages (Sheward et al., 2018; Williams et al., 2018). Whereas in a paradigm of coevolution, one might expect that novel epitopes introduced by a superinfection would spur the development of existing HIV-targeting Ab lineages to evolve innovations to regain affinity. Similarly, in the context of malaria infection, repeated immunizations with a complex malaria antigen promote the activation of *de novo* naïve Ab lineages, rather than the evolution of already existing malaria-targeting Abs (Murugan et al., 2018).

A better understanding of the interaction between HIV and the AbR over time could shed light on how HIV immunity is achieved. However, longitudinal sequence studies of HIV infections tend to either focus on HIV (Feder, Kryazhimskiy, & Plotkin, 2014; Fischer et al., 2010; Henn et al., 2012; Shankarappa et al., 1999; Zanini et al., 2015) or the AbR (Hoehn et al., 2015; Nourmohammad, Otwinowski, Luksza, Mora, & Walczak, 2018; Xueling Wu et al., 2015), but not both. To our knowledge, only three studies have deeply sequenced the AbR along with the autologous HIV population, but each of these studies consisted of a single individual with relatively few time-points (Bhiman et al., 2015; Landais et al., 2017; Liao et al., 2013). In this study, we ameliorate this dearth of data, and describe in unprecedented detail the genetic interaction between these putatively coevolving populations in 10 well-characterized HIV-infected individuals.

## Results

### Sequencing HIV *env* and IGH

We collected a total of 119 cryopreserved peripheral blood samples from the OPTIONS cohort at the University of California, San Francisco (UCSF). The samples all originated from male participants aged 25-48 years old at the estimated time of infection in San Francisco, and each participant had 10-20 longitudinal samples (Table 1, and Figure S1). The samples were all collected prior to administration of ART, with the exception of the last time-point of participants 1, 2, and 5. We chose to deeply sequence the C2-V3 region of the *env* gene because of its rich history of interactions with Abs, as evidenced by the HIV epitope map from Los Alamos National Labs (LANL) (Hatada et al., 2010; Ringe et al., 2012; Yusim et al., 2016). Of the 116 ART-naïve samples, we were unable to successfully amplify C2-V3 from 12. In 11 of these cases, low viral load was the presumed cause for lack of amplification, but the 9^th^ time-point of participant 6 was unsuccessful despite high viral load (Figure 1, S2). We were also unable to amplify C2-V3 from the 3 ART-exposed samples. While viral load measurements were not available for these samples, the presumed cause for our inability to amplify C2-V3 was low viral load, given their ART status. The initial *env* sequencing depth ranged from 3,771-101,831 reads per sample, and after cleaning the data with several quality control (QC) steps, this ranged from 2,276-56,914 reads per sample (Figure S3).

**Table 1.** Participant demographics.

| ID | Age* | Date* | Gender | Ethnicity | Exposure | Num. Samples |
| --- | --- | --- | --- | --- | --- | --- |
| 1 | 30 | 6/7/98 | Male | White/European American | MSM | 20 |
| 2 | 25 | 2/17/99 | Male | Asian | MSM | 17 |
| 3 | 32 | 7/4/01 | Male | Hispanic/Latino | MSM | 10 |
| 4 | 44 | 8/8/01 | Male | White/European American | MSM | 10 |
| 5 | 34 | 7/20/03 | Male | Hispanic/Latino | Unknown | 11 |
| 6 | 25 | 1/18/05 | Male | White/European American | MSM | 10 |
| 7 | 35 | 6/6/05 | Male | White/European American | MSM | 10 |
| 8 | 48 | 9/3/08 | Male | White/European American | MSM | 10 |
| 9 | 33 | 2/2/09 | Male | Asian | MSM | 10 |
| 10 | 31 | 4/3/09 | Male | White/European American | MSM | 10 |

MSM – men who have sex with men.
\* at estimated time of infection

**Figure 1.**
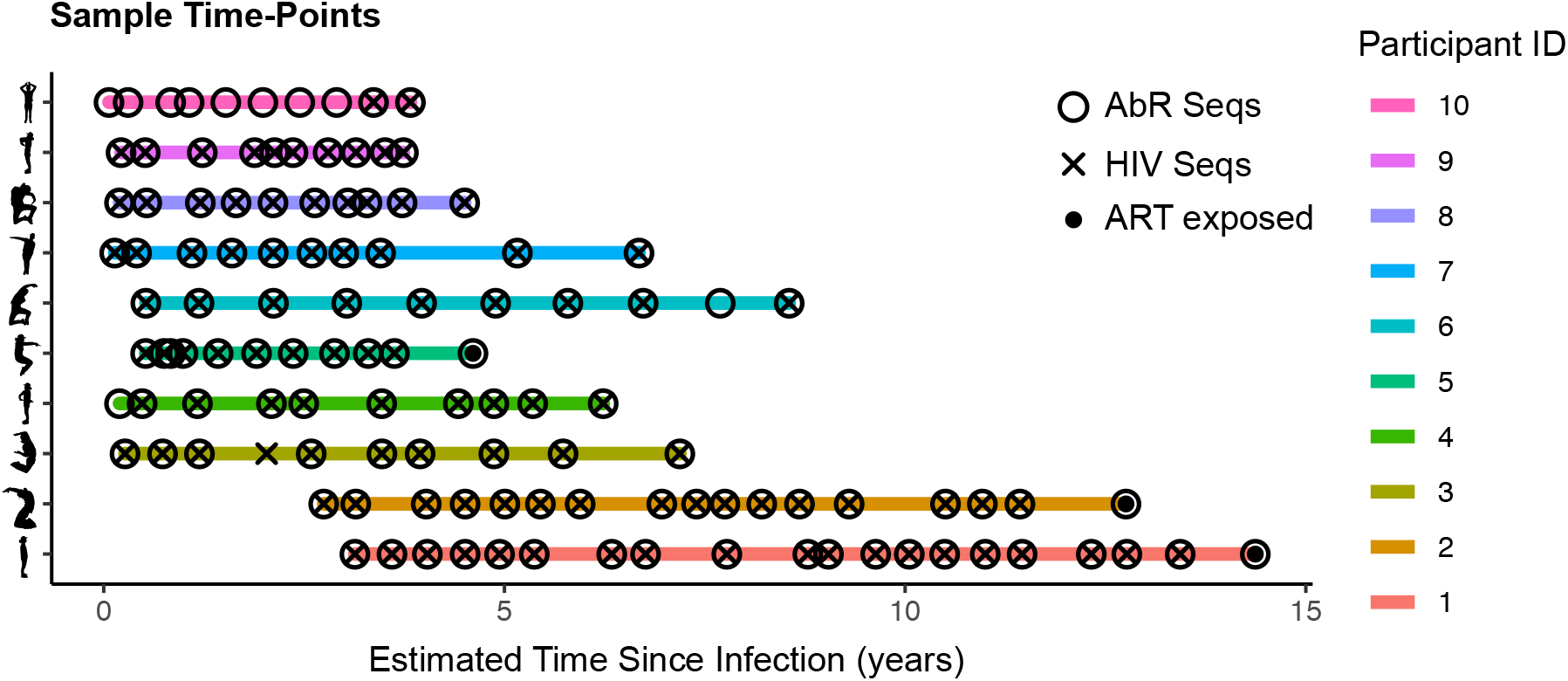
Sampled time-points. Depicts the time since infection for each sample, in each of the participants. Open circles indicate that the AbR was successfully sequenced, crosses indicate that HIV was successfully sequenced, and solid circles indicate that the participant was on ART at this time.

We also deeply sequenced the variable region of the immunoglobulin heavy chain locus (IGH), the product of which we refer to as the antibody repertoire (AbR). The AbR was successfully sequenced in all 119 samples, with the exception of the fourth time-point of participant 3. Sequence data was generated for this sample, but it exhibited an abrupt clonal expansion of a magnitude that was a clear outlier for participant 3, and not seen in any other sample, so it was discarded (Figure S4, and 1). Initial AbR sequencing depth ranged from 669,331-2,669,662 reads per sample, and after QC this ranged from 160,291-552,479 reads per sample (Figure S5).

### Characterizing the HIV population and AbR over time

In order to quantify the broad attributes of the AbR and HIV populations over time, we calculated a variety of summary statistics that characterized the genetic diversity, divergence, selection, and abundance for each of the populations. Selection pressure (positive or negative) in HIV was estimated as *dN*/*dS* (Nei & Gojobori, 1986), and in the AbR was estimated using the ∑ values from BASELINe (Yaari, Uduman, & Kleinstein, 2012). As others have reported (Maldarelli et al., 2013; Zanini et al., 2015), we found that HIV diversity, divergence, and selection (*dN*/*dS*) all tend to increase with time since estimated date of infection, with large perturbations over smaller time-scales (Figures 2, S6, and S7). Of note, the high viral load of the first time-point of participant 7 suggests that the acute viremia phase of early HIV infection was captured. Additionally, participant 10 had very low viral load for the first 8 time-points, which explains why amplification of HIV *env* was unsuccessful for these samples. Participant 6 exhibited strong evidence for a super-infection occurring between the 1^st^ and 2^nd^ time-points (Figure S8). Super infections are not uncommon with HIV (Redd, Quinn, & Tobian, 2013), and will cause a sudden injection of ‘artificial’ genetic divergence relative to the initial infecting virus. Thus, we accounted for this superinfection when calculating divergence summary statistics for participant 6 by comparing each sequence to both the initial infection, as well as the subsequent superinfection (see Methods).

**Figure 2.**
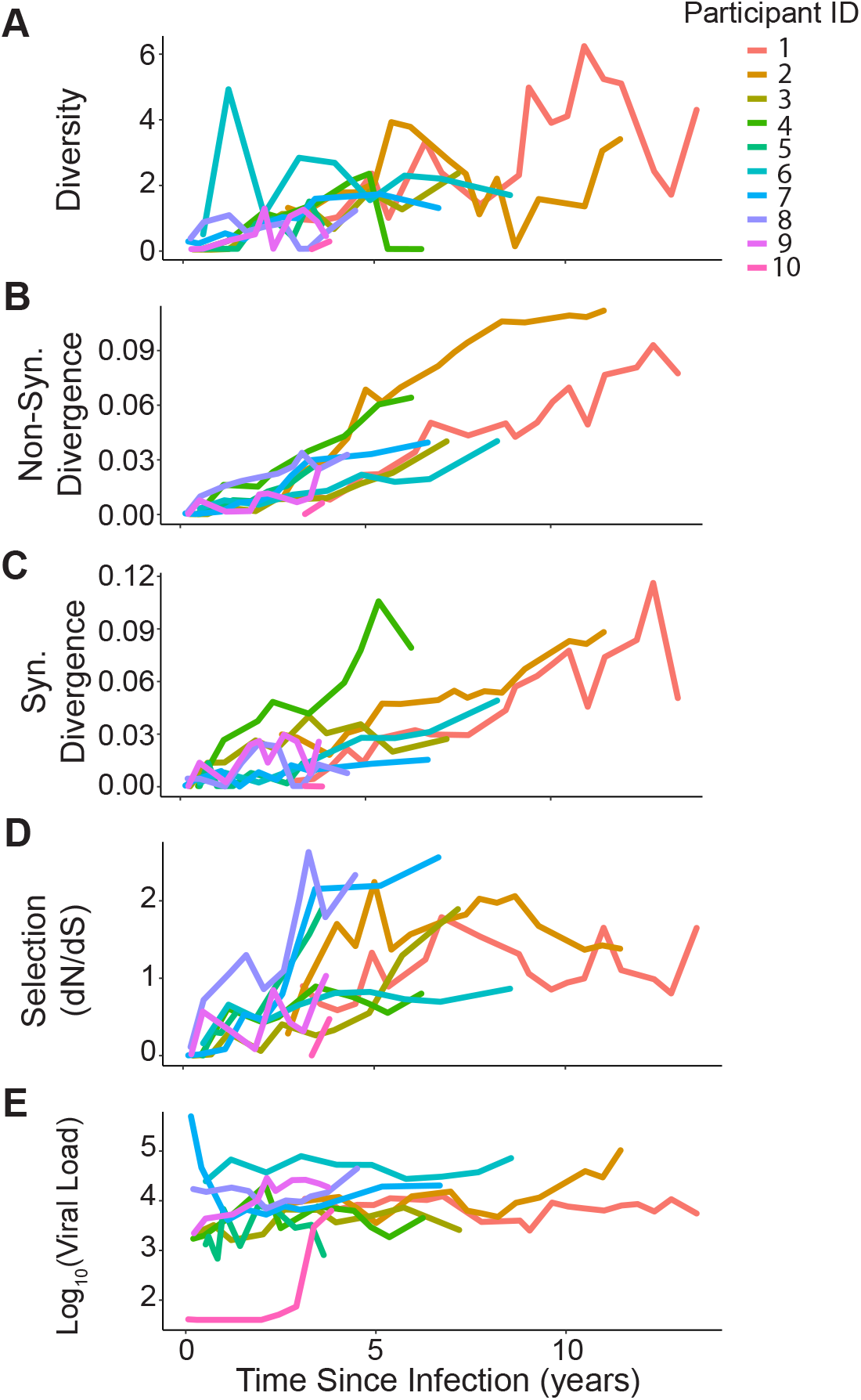
HIV summary statistic trajectories for each participant. Each line shows the trajectory of a given summary statistic, in a given participant over time. Each participant has a unique color. (A) Diversity. (B) Nonsynonymous divergence. (C) Synonymous divergence. (D) Selection. (E) Viral load.

The trajectories of the AbR summary statistics did not show any obvious stereotyped pattern across participants (Figures 3, S9, and S10). However, there were a couple data points that suggested interactions with the HIV population: the second time-point of participant 7 (0.41 years post infection) showed a large increase in selection (in both the framework regions, FWR, and complementarity determining regions, CDR), which could be a response to the initial viremia in the prior time-point; and the ninth time-point of participant 10 (3.37 years post infection) also showed a large increase in selection with a concomitant drop in diversity, which could have been a response to the large increase in viral load at that time-point.

**Figure 3.**
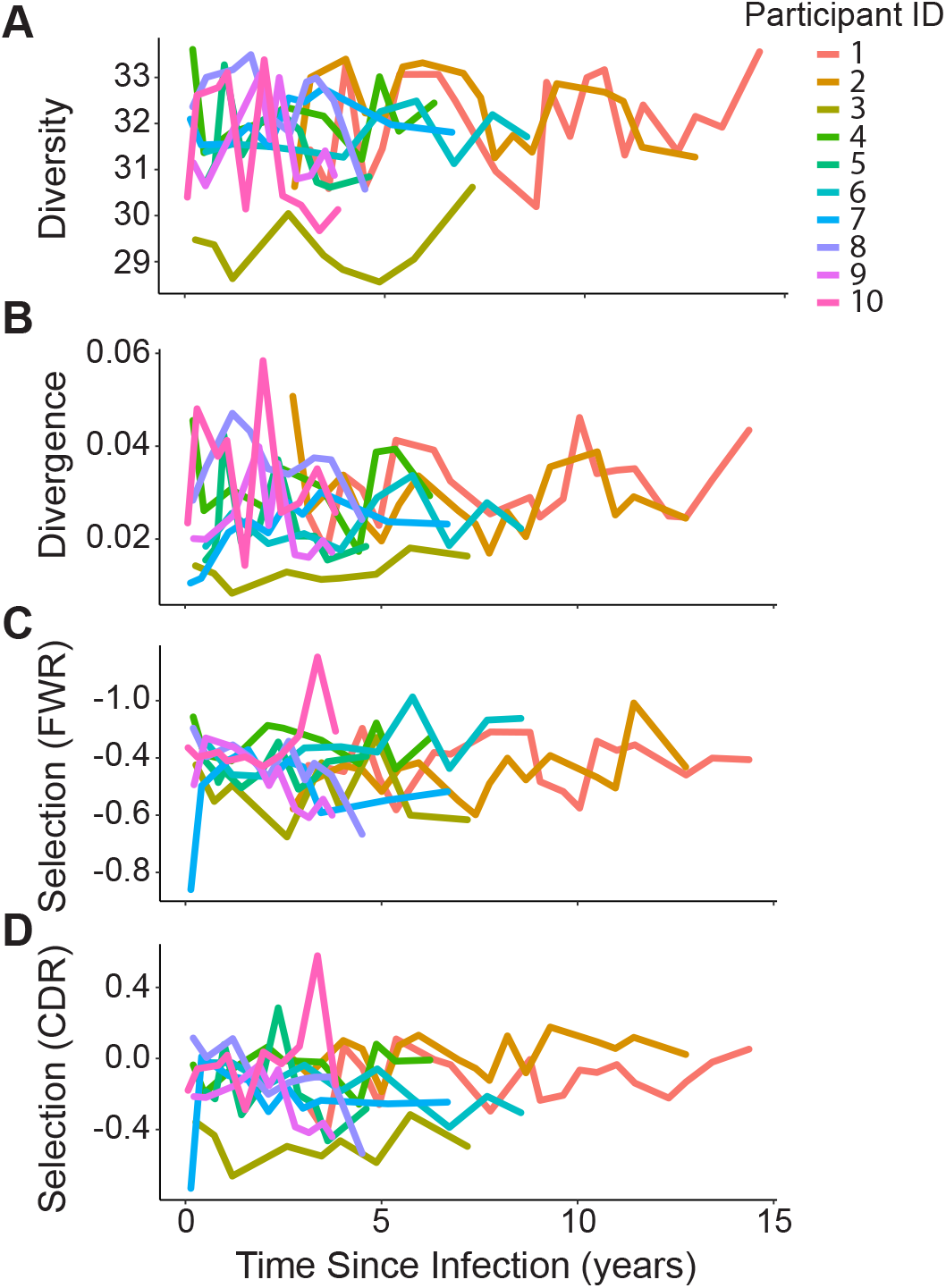
AbR summary statistic trajectories for each participant. Each line shows the trajectory of a given summary statistic, in a given participant over time. Each participant has a unique color. (A) Diversity. (B) Divergence. (C) Selection in the framework region (FWR). (D) Selection in the complementarity determining region (CDR).

To increase the accessibility of these data, we created a web application, available at https://ab-hiv-coevolution.github.io/HIV_AB_CoEvo/, for others to investigate this new dataset interactively. It allows users to explore the data by individual patient or all patients across various summary statistics including genetic diversity, divergence, selection, and abundance for HIV and/or the AbR.

### Testing for whole-population level interactions

In order to gauge the effect HIV has on the AbRs of participants, we first pooled all the data across participants, and used a regression framework to test if any of the AbR summary statistics were significantly correlated with that of the HIV population (while controlling for participant-specific effects, see Methods). We performed this test in a pairwise fashion on all AbR summary statistics against all HIV summary statistics. Similar to Hoehn’s work (Hoehn et al., 2015), we found no significant correlation between AbR diversity and viral load, yet we did find a modest, statistically significant correlation between AbR and HIV diversity (p= 0.02, Figure 4 A). While this association was marginally significant—indeed, it ceases to be significant after controlling for multiple tests—we found any association at the whole-population level surprising, and thus, worth reporting.

**Figure 4.**
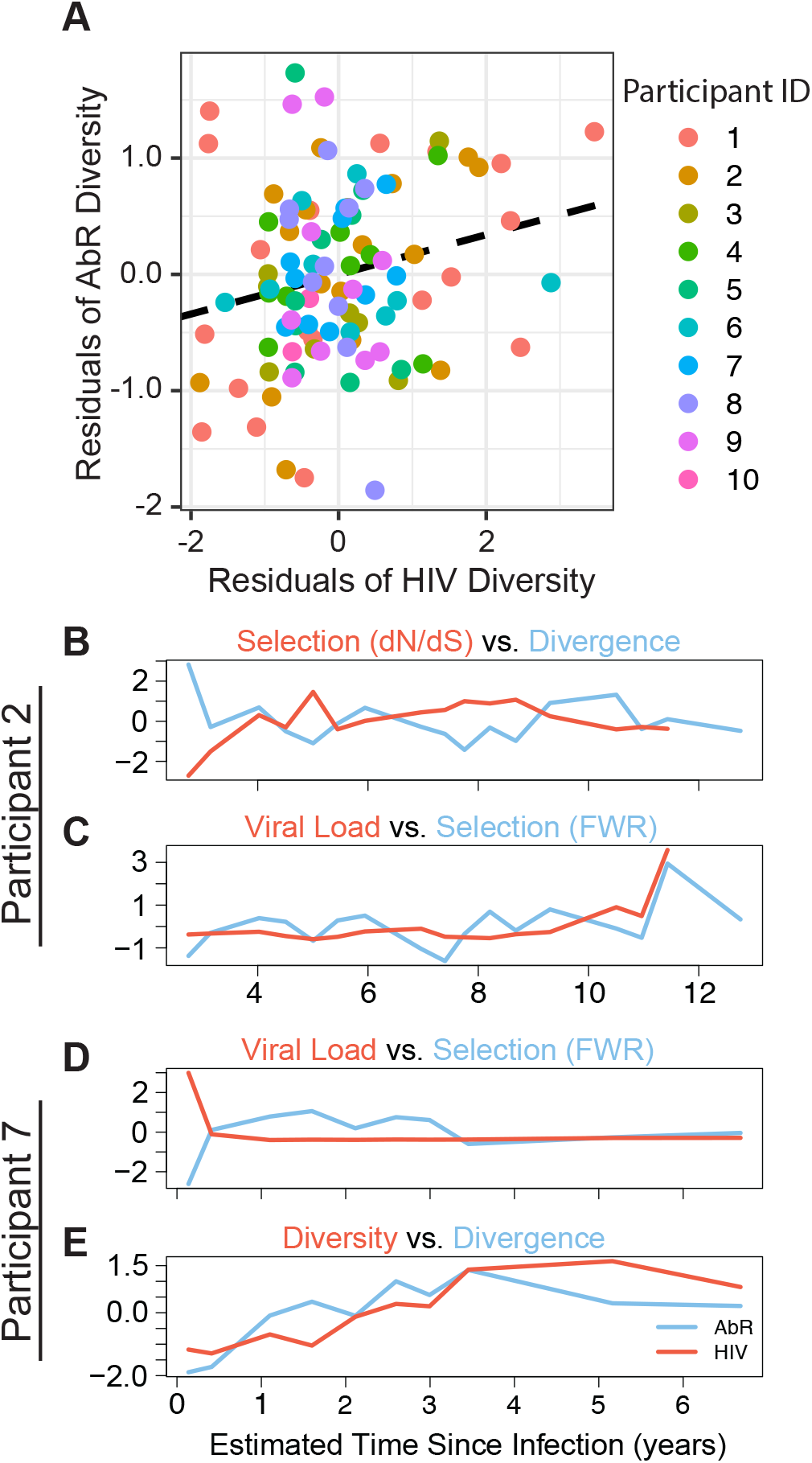
Whole population level associations between AbR and HIV summary statistics. (A) Scatter plot showing positive correlation between HIV diversity (x-axis) and AbR diversity (y-axis). Each point represents a sample with diversity values from both the AbR and HIV sequence data. The axes show diversity values after participant specific effects have been regressed out. Dashed line shows positive relationship between AbR and HIV diversity, as given by our linear regression (see Methods). (B-E) Shows associations between summary statistic trajectories in AbR (blue) and HIV (red) at the individual participant level. (B) HIV selection with AbR Divergence, in participant 2. (C) Viral load with AbR FWR-selection, in participant 2. (D) Viral load with AbR FWR-selection, in participant 7. (E) HIV diversity with AbR divergence, in participant 7.

It is possible that some participants’ AbRs interact with their autologous HIV population more than others, thus we also tested for interactions between summary statistics on a participant-by-participant approach. Because the number of data points per participant is relatively low, we opted to use a permutation based test in order to accurately estimate type 1 error (Ernst, 2004) (see Methods). In participant 2, we found that selection (*dN*/*dS*) and viral load in the HIV population were associated with divergence and selection (FWR) in the AbR, respectively (p=0.009 and p=0.0425, Figure 4 B, and C). We also found that in participant 7, viral load and diversity in the HIV population were associated with selection (FWR) and divergence in the AbR, respectively (p=0.040 and p=0.040, Figure 4 D, and E). All of these associations were positive correlations, with the exception of HIV selection (*dN*/*dS*) and AbR divergence in participant 2, which was anticorrelated.

Together, these data raise the possibility that the effect that HIV has on the AbR (and vice versa) is large enough to be observable at the intra-participant, whole population level, although more samples would be needed to make definitive statements about this.

### Identifying partitions of the AbR that interact with HIV

The AbR is an exceedingly complex population consisting of a myriad of Ab lineages capable of simultaneously binding and neutralizing a countless number of antigenic targets. To identify specific parts of the AbR that may be interacting with HIV, we first partitioned the AbR across time based on the germline identity of each sequence’s V and J gene segments (Figure S11), and then tested each AbR partition for evidence of interactions with the autologous HIV population using similar summary statistics as the overall population (see Methods). Using an analogous permutation-based test as was used when comparing the overall populations, we found significant associations between AbR-partition trajectories and HIV trajectories in participants 3, 7, and 8 (Figures S12, S13, and 5). Of these associations, AbR-partition frequency tended to be most often associated with viral load. For example, participant 7 had a distinct viral load trajectory— presumably due to acute viremia—and the frequency trajectory of the IGHV4-31:IGHJ5 AbR partition was positively associated with the unique shape of this trajectory (while the diversity of this AbR partition was negatively associated with viral load). This suggests a clonal expansion occurred in this AbR-partition in response to HIV, which caused an increase in the partition’s frequency with a concomitant drop in diversity. Similarly, in participant 8, the frequency trajectory of two AbR partitions with the same V gene segment (IGHV6-1:IGHJ5 and IGHV6-1:IGHJ4) were positively associated with viral load, suggesting that the IGHV6-1 gene segment in this participant may have had a predisposition to targeting HIV. In participant 3, the frequency trajectory of the IGHV3-30:IGHJ3 partition was negatively associated with both non-synonymous divergence and selection (*dN*/*dS*) in the HIV population, suggesting that escape mutations in HIV caused a drop in frequency of the interacting AbR partition.

### Validating the HIV-associated AbR partitions

In order to establish that our permutation-based test is in fact identifying AbR partitions that had a biological response to HIV infection and was not the result of random noise in the data, we sought to compare our results to previous findings in the literature. We first used Fisher’s method to compile all the results of our permutation-based test into a single score for each V gene segment (see Methods), and then used the database of HIV bnAbs from bNAber (Eroshkin et al., 2014) to compare the incidence of known HIV-binding V gene segments in the literature, to how well these V gene segments score in our test (Table S1). We found that V gene segments that had been shown to bind HIV in the literature dataset were associated with significantly higher scores in our permutation-based test (p=1.52e-3, Mann-Whitney U test, Figure S14 A, and C). There are two reasonable explanations for this: i) these V gene segments have a predisposition to bind HIV (i.e. a ‘public’ or convergent response (Setliff et al., 2018)), or ii) These V gene segments have a predisposition to bind anything (due to high endogenous expression, having broad affinity for viruses, or some other unknown reason). In order to differentiate between these two possibilities we performed a similar test except instead of comparing our results to Abs known to target HIV, we compared our results to a literature dataset that we previously compiled of Abs that have been shown to bind to influenza (Strauli & Hernandez, 2016) (Table S1). Similar to the HIV literature dataset, we found that V genes that were well represented in the influenza literature dataset also tended to score highly in our permutation-based test (p=7.28e-4, Mann-Whitney U test, Figure S14 B, and D). Therefore, while we are likely identifying a biological response to HIV, the response may not be specific to HIV.

### Testing for coevolution

Coevolution between HIV and a handful of well-known bnAbs has been extensively reported (Liao et al., 2013; Rantalainen et al., 2018) and reviewed (Bonsignori, Liao, et al., 2017). Coevolution provides an intellectually compelling explanation for the development of bnAbs against HIV, however, examples tend to be anecdotal and qualitative (likely due to small sample sizes). While we cannot be sure that bnAbs exist in our data, we sought to test if coevolution is a predominant driver of HIV-targeting Ab development generally. We tested for genetic signals of coevolution in our data by first dividing the AbR data of each participant into time-course lineages of Abs (Figure 6, S15, and S16). We then used MAFFT (Katoh, Misawa, Kuma, & Miyata, 2002) to create a multiple sequence alignment (MSA) of each Ab lineage, and compare each of these Ab lineage MSAs with a representative MSA of the HIV population overtime using a mutual information (MI) statistic (Brandman, Brandman, & Pande, 2012; Gloor, Martin, Wahl, & Dunn, 2005; Marino Buslje, Teppa, Di Doménico, Delfino, & Nielsen, 2010; Simonetti, Teppa, Chernomoretz, Nielsen, & Marino Buslje, 2013). Importantly, we reduced the complexity of the amino acid code to a code of ‘change’ or ‘no-change’ prior to calculating MI (see Methods).

**Figure 5.**
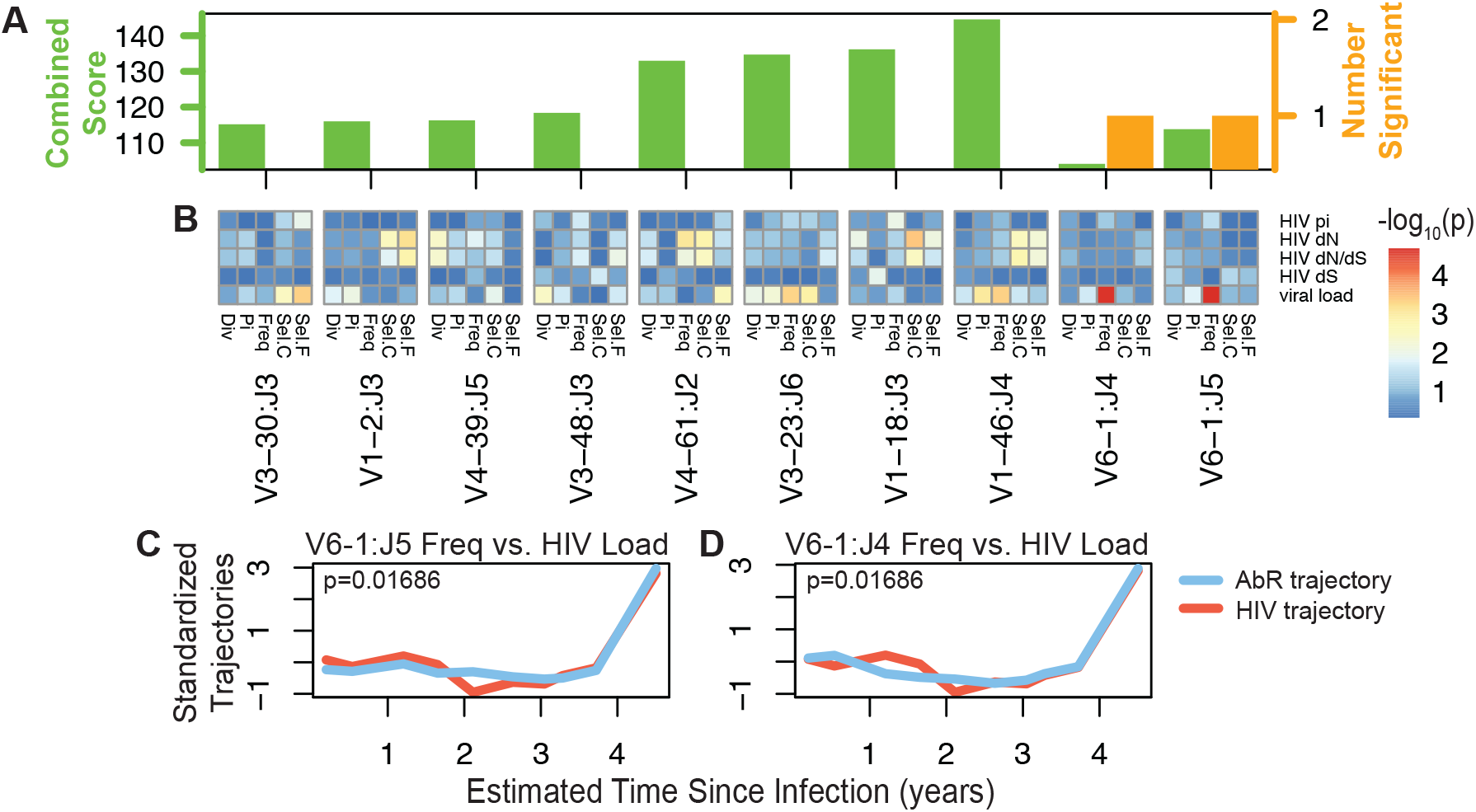
Results of permutation-based test to identify HIV-associated AbR partitions in participant 8. (A) A barplot showing the combined score from the permutation-based test (left axis), and the number of significant associations (right axis) for the top 10 AbR partitions. AbR partitions were sorted first by the number of significant associations, then by their combined score from the permutation-based test (ascending, left-right). (B) Heatmaps depicting the significance (−log_10_(p-value)) for each permutation test within each of the top 10 AbR partitions. Columns correspond to summary statistics of the AbR partitions: Div=divergence, Pi=diversity, Freq=relative frequency, Sel.C=CDR selection, and Sel.F=FWR selection. Rows correspond to summary statistics of the HIV population: HIV pi=diversity, HIV dN=nonsynonymous divergence, HIV dS=synonymous divergence, HIV dN/dS=selection, and viral load is self-explanatory. The color of each element in the heatmaps shows the significance of the association between a given AbR partition summary statistic trajectory with a given HIV population summary statistic trajectory. (C, D) Shows the AbR partition (blue) and HIV population (red) trajectories that were significantly associated. (C) Frequency trajectory of the IGHV6-1:IGHJ5 AbR partition with the viral load trajectory of the HIV population. (D) Frequency trajectory of the IGHV6-1:IGHJ4 AbR partition with the viral load trajectory of the HIV population.

**Figure 6.**
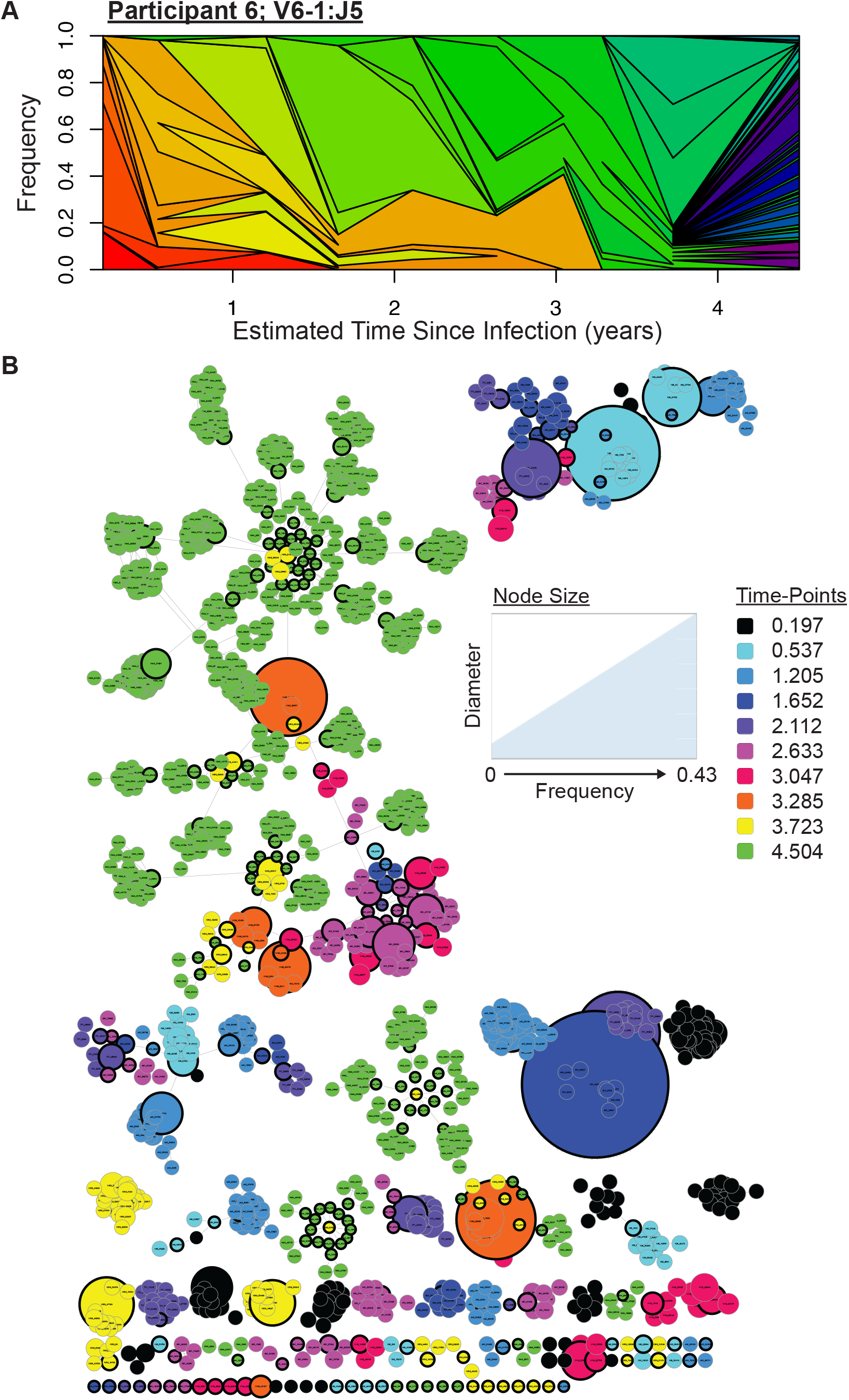
AbR lineage structure. Illustrates the lineages in AbR partition IGHV6-1:IGHJ5, participant 8. (A) Muller plot depicting the relative frequency of each lineage within the partition. Each color represents a lineage. If a new lineage arises within the bounds of a preexisting lineage, then the new lineage is a daughter of the preexisting, parent lineage. Lineages that began earlier in the time-course have colors closer to the red side of the spectrum, while lineages that began later in the time-course have colors closer to violet. Only lineages that exceeded 0.0001 frequency are included in the plot. (B) Network plot. Each node represents a unique sequence. Nodes with thicker black borders are the ‘representative sequence’ of their sequence-cluster. Nodes of the same color (i.e. same time-point) are linked with an edge if they were in the same cluster. Nodes that have different colors are linked with an edge if they were assigned to the same lineage. This inter-time-point linking only occurs between ‘representative sequences’. Taken together, each isolated grouping of nodes shows a family of related lineages.

If coevolution is a common characteristic of HIV-targeting Ab lineages, then we might expect that the HIV-associated AbR partitions that we previously identified will have abnormally high MI values. Thus, we compared the mean MI values of the lineages within all of the HIV-associated AbR partitions (IGHV3-30:IGHJ3 of participant 3, IGHV2-70:IGHJ6 of participant 7, IGHV3-15:IGHJ4 of participant 7, IGHV4-31:IGHJ5 of participant 7, IGHV6-1:IGHJ4 of participant 8, IGHV6-1:IGHJ5 of participant 8) to the distribution of mean MI values from the rest of the lineages (Figure 7 A). In addition to mean MI, we also compared the length of lineages (i.e. the number of time-points for which a lineage is present in the data), because coevolving lineages might be expected to persist in the AbR longer than non-coevolving lineages. We found no evidence of coevolution in the HIV-associated AbR partitions, with the exception of a single lineage in IGHV6-1:IGHJ5 of participant 8, which had a mean MI value that was in the 99.55^th^ percentile relative to all the Ab lineages in non-HIV-associated AbR partitions. We found that this result persisted when comparing to a simulated null distribution that controls for uncertainty when assigning lineages across time-points (Figure 7 B).

**Figure 7.**
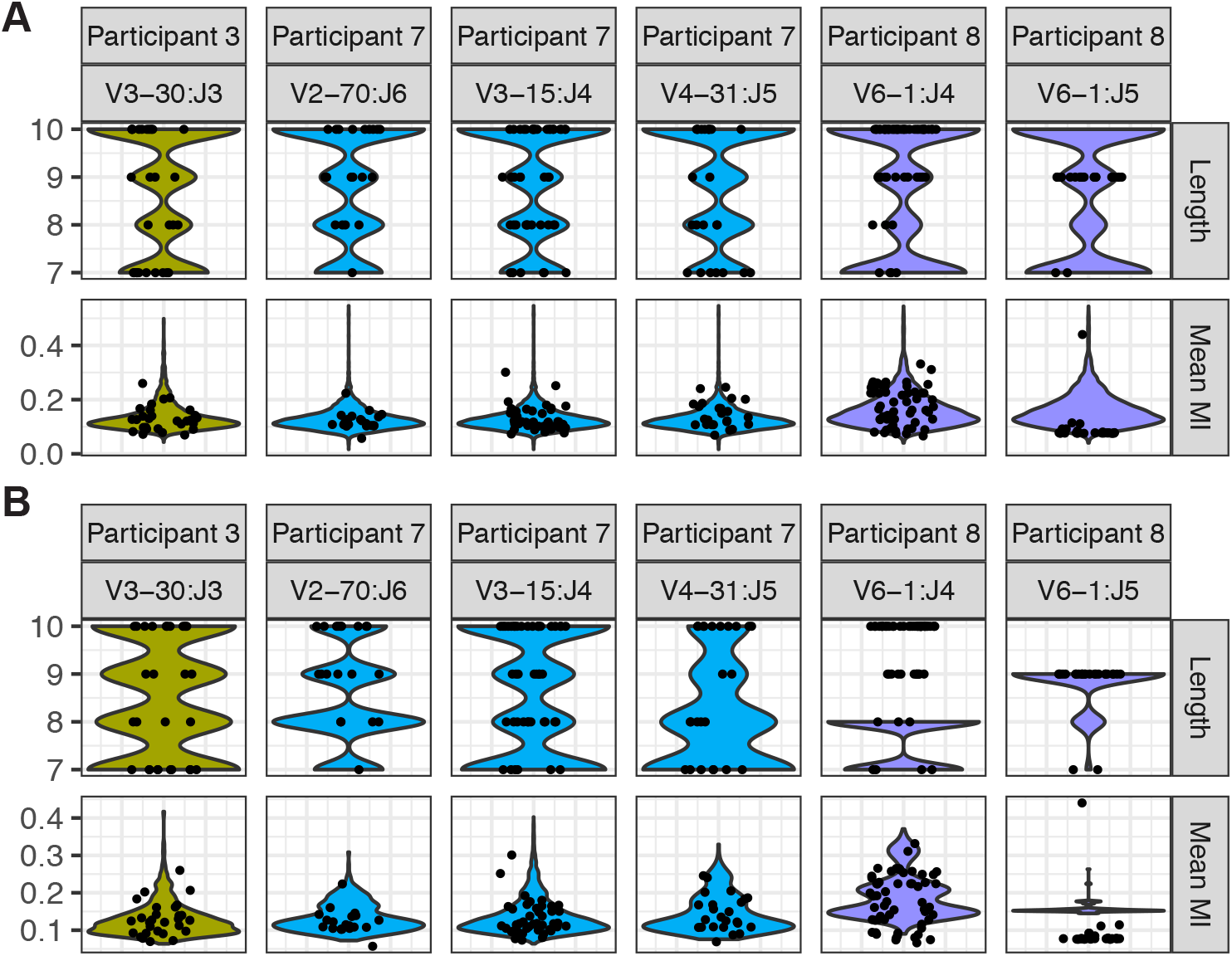
Coevolution test for individual lineages. Each black dot corresponds to a value for a specific lineage within an HIV-associated AbR partition, and are jittered across the x-axis. The identifying information for the specific HIV-associated partition is given by the column labels for the plots (ex. The values for the lineages within the IGHV3-30:IGHJ3 AbR partition of participant 3 are found in the left-most column). The colored violin plots behind the points represent a given null-comparison for the points, where color corresponds to participant. The top row of plots shows the values and null distributions for the length of lineages (i.e. how many time-points they were present). The bottom row shows the values and null distributions for the mean MI of lineages. (A) The null distributions are made up of the values from the lineages in AbR partitions that were not HIV-associated. (B) The null distributions are made up of the values from the null simulation for each given AbR partition.

Lastly, we test for a global coevolutionary signal, agnostic to whether or not a lineage belongs to an HIV-associated partition. To do this we gathered the mean MI values across all of the observed lineages within a participant, and then compare this distribution to that of the mean MI values from the simulated null lineages (Figure 8). If Ab/HIV coevolution were taking place on a large scale, we would expect to see a shift towards higher MI values in the observed distribution relative to the simulated null. However, we saw no evidence of this, and instead saw that, if anything, the simulated null lineages tend to have higher MI values. This suggests that if coevolution is taking place at all, it is either a weak force or rare in these participants.

**Figure 8.**
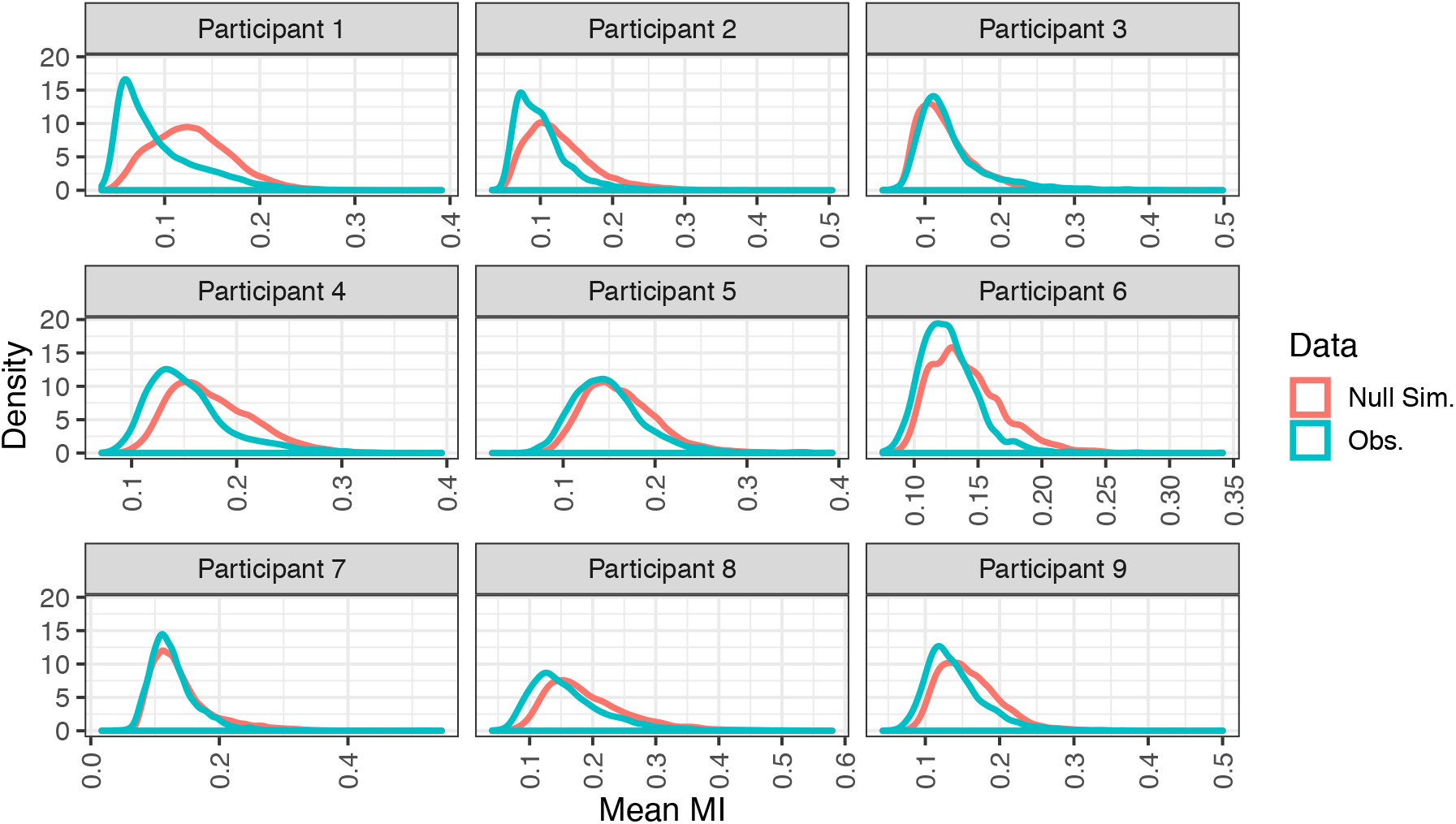
Global tests for coevolution. Each line gives the distribution of mean MI values for a given participant. Turquoise colored lines correspond to observed lineages (i.e. lineages inferred from the data), and salmon colored lines correspond to lineages from the null simulations. Participant 10 is omitted because of limited HIV sequence data.

## Discussion

In this study we have created a large dataset of Ab sequences and HIV sequences from 119 longitudinal samples. This is, to our knowledge, the largest dataset of its kind. The HIV literature currently encompasses an abundance of AbR sequence datasets from HIV+ individuals. However, these datasets primarily originate from the amplification of a particular Ab lineage that was known to contain an HIV-bnAb, prior to deep sequencing. These ‘biased’ AbR datasets are useful to home in on the development of a particularly interesting Ab lineage. However, we argue that it is equally important to understand the AbR response to HIV infection on a global/systems scale for the following reasons: i) It is just as important to understand why an HIV-targeting Ab lineage failed to develop broad neutralization ability, as it is to understand why a lineage succeeded in developing it. ii) There is increasing evidence that non-neutralizing Ab lineages play important roles in viral pathogenesis through Fc-mediated functions (e.g. antibody-dependent cellular cytotoxicity (Horwitz et al., 2017; Mayr, Decoville, et al., 2017; Mayr, Su, & Moog, 2017). iii) It is quite possible that a significant proportion of humans that develop broad immunity to future HIV infections, do so in a polyclonal manner (Williams et al., 2018), where broad neutralization depth against HIV is achieved via the cooperative action of many Ab lineages, each simultaneously targeting different epitopes, or different versions of the same epitope. iv) The population dynamics of HIV-bnAbs (in addition to HIV-binding Abs in general) are poorly understood. For example, do these Abs persist at high or low frequency in the greater population? How does this frequency change over time? What type of selection drives their development (ex: positive, negative, balancing, etc.)? These types of questions are difficult, if not impossible to answer without understanding the larger population context for which these Ab lineages exist. In this study, we have taken the preliminary steps towards addressing these types of questions. Specifically, we have developed a statistical approach to identify the partitions of the AbR that are likely responding to HIV. Once this has been established, questions like those enumerated above can be answered. Further, we hope that the sequence datasets we have created here will provide a useful resource for others with similar lines of inquiry.

While (Hoehn et al., 2015) found no correlation between AbR diversity and viral load in their data, they did not have the means to address other characteristics of the HIV population, as they did not include HIV sequence data in their analyses. However, they did find that AbR diversity was lower in HIV+ individuals than healthy controls. This suggested that HIV may have a broad effect on the AbR, yet the details of this effect remained unclear. We have presented a small positive correlation between overall AbR and HIV diversity across all our samples. A possible scenario that would explain this observation is one where AbR diversity is decreased due to clonal expansions in Ab lineages that target HIV, which in turn causes a decrease in HIV diversity, due to positive selection for escape mutations. Once the HIV population has escaped, its diversity will return, and diversity in the AbR will also return because the previous clonal expansion will have vanished due to its target having escaped. However, we stress that this correlation had nominal significance and should be treated cautiously. The AbR is an especially complicated population that is capable of simultaneously responding to countless antigens, thus even a fleeting correlation with HIV at the whole-population level may be worthy of further follow up studies.

A key first step towards illuminating the global interaction between the AbR and an HIV infection is to be able to identify the subset of the AbR that is responding to HIV. Similar to our previous work in the context of influenza vaccination (Strauli & Hernandez, 2016), we leveraged the time-series nature of our dataset to identify partitions of the AbR that seem to be associated with HIV. We purposefully made no prior assumptions about what types of interactions we might find. For example, the common narrative of HIV-targeting Ab lineages is that they are under intense positive selection. Thus, one might have the expectation that Ab selection will be positively correlated with HIV selection. However, it is also possible that an HIV-targeting Ab lineage is under intense negative selection, where there is a preference for amino acids *not* to change so as to not ablate their binding ability, or perhaps to not change a strict structural conformation that is required to access an epitope. In this case, one might expect Ab selection to be negatively correlated with HIV selection. We therefore compared all AbR summary statistics to all HIV summary statistics. This all-by-all comparison gave us the privilege of an unbiased approach, yet greatly increased the number of tests that were performed, and hence the severity of our multiple-tests correction. As such, we were only able to identify a handful of AbR partitions that were significantly associated with HIV. This suggests that long-term interactions between Ab lineages and HIV are rare, and that Ab/HIV interactions may be of a more transitory nature, where an antibody binds to HIV, then HIV escapes, and then another unrelated Ab binds to the escape mutant, and so on. Another possibility is that our test was simply underpowered and had many false negatives. One way to ameliorate this would be to first filter AbR partitions based on some statistic (e.g. divergence) and then test for associations using a different, orthogonal statistic (e.g. diversity). Further, there remains a great deal of powerful analyses that could be done with the HIV sequence data. In principle, one could divide the HIV population into lineages and test each HIV lineage against each AbR partition. This would increase the number of tests, but could also illuminate interactions that would be otherwise hidden.

Lastly, we tested for a coevolutionary signal in our data. Coevolution in sequence data is notoriously hard to establish (Avila-Herrera & Pollard, 2015), and to our knowledge, reports of HIV/Ab coevolution to date have been universally qualitative, with little or no statistical analyses (Bhiman et al., 2015; Bonsignori, Kreider, et al., 2017; Bonsignori et al., 2016; Nicole A. Doria-Rose et al., 2014; Gao et al., 2014; Landais et al., 2017; Liao et al., 2013; MacLeod et al., 2016; Rantalainen et al., 2018). When it comes to claims of coevolution, there are two sources of uncertainty that we have attempted to account for in this study. i) When both the Ab lineage and the putative HIV epitope are under positive selection, it is very easy for mutations to be correlated by chance rather than by coevolution. ii) There is a huge amount of uncertainty when assigning Abs to lineages, especially when trying to link a given Ab sequence to other Ab sequences that existed months-years prior. AbRs are incredibly dynamic populations with high turnover, and high mutation rates. In a population such as this, where *de novo* lineages are continuously being added, it is important for one to account for the possibility that two similar Ab sequences—even if strikingly similar—may not be of the same lineage. By creating a simulated null dataset from shuffled Ab lineages, we were able to create a null distribution of MI values that took both of these confounders into account. After doing this we found no global signal for coevolution, yet we did find one isolated Ab lineage in participant 8 that showed compelling evidence for it. This suggests that while coevolution between Ab lineages and HIV is possible, it is likely exceedingly rare and/or hard to detect. Given that other sequence datasets of the AbR in the context of HIV infection have about the same or fewer time-points than the participants in our dataset, we suspect that claims of coevolution in these data would be equally hard to make.

Coevolution has been responsible for some of the most remarkable phenotypes known (e.g. the cheetah’s speed, a flower’s beauty, the strangeness of genitalia (Brennan & Prum, 2015)), yet it remains unclear as to how much of a role it plays in the development of HIV-targeting Abs. This has important implications for vaccine strategies. If coevolution is the predominant force in the development of HIV-bnAbs, then a vaccine regimen that mimics the natural evolution of the HIV epitope *in vivo* would be desired, as this would recapitulate the coevolutionary process leading to increased neutralization breadth. However, it is also possible that HIV-bnAbs occur as rare, random events. Examples of this include unlikely V(D)J recombination events whereby a naïve Ab lineage is created that happens to have a predisposition to target a conserved HIV epitope, or a (typically diverged) Ab lineage ‘stumbling’ upon broad neutralization breadth by chance. In this case, one might desire a vaccine regimen that has a very diverse array of HIV epitopes so as to maximize the chances that these rare events occur. This approach is supported by the fact that neutralization breadth is positively correlated with viral load and HIV diversity (N. A. Doria-Rose et al., 2010; Landais et al., 2016; Piantadosi et al., 2009). An interesting future study would be to use mathematical modeling (Nourmohammad, Otwinowski, & Plotkin, 2016), or simulation frameworks (Murugan et al., 2018) that include the introduction of novel naïve Ab lineages into the population, to gain a better understanding of which evolutionary parameters (e.g. population size, mutation rate, selection strength, population diversity, etc.) in the HIV and AbR populations promote coevolution, and which do not. We hope that this study has provided a sound example of how to go about formally testing for coevolution in order to differentiate between these two possibilities.

## Methods

### Participant selection and sample processing

Samples were obtained from the OPTIONS cohort at UCSF. All participants provided written informed consent, and the study was approved by the UCSF Committee on Human Research. Our sole criterion for selecting participants from this cohort was to find those with the greatest number of samples available prior to the administration of ART. All the participants in this study were men who contracted HIV via sexual transmission, with the exception of participant 5, who became infected by unknown means (Table 1). Each peripheral blood sample was divided into plasma and peripheral blood mononuclear cells (PBMCs) by density gradient centrifugation using Ficoll-hypaque. After separation, PBMCs and plasma were aliquoted in cryopreservation media, and cryo-preserved in a specimen repository. Plasma viral load was measured at each participant visit.

### Viral load

In early samples (prior to ~2009), a combination of a branched DNA assay and an ultra-sensitive PCR assay from Roche were used to measure HIV load. In later samples (post ~2009), the Abbott RealTime HIV-1 Viral Load assay was employed to measure load.

### Estimated time of infection

The time of initial infection was estimated using the following criteria: i) If a participant first presents with detectable viral load, but negative enzyme immunoassay (EIA) or western blot, and then presents a positive western blot in the following visit, then the estimated time of infection is given by 24 days prior to the first visit. ii) If a participant first presents with an indeterminate western blot, and then a positive western blot following repeat testing, then the estimated time of infection is given by 24 days prior to this first test. iii) If participant first presents with a positive western blot, and has a documented negative HIV test result within at most 180 days prior to first test, then the estimated time of infection is calculated as 24 days prior to the midpoint between the first positive test and prior negative test. We note that first visit here corresponds to the first visit in the OPTIONS study at UCSF, and not the first sample in our study.

### C2V3 amplification and ultra-deep sequencing

HIV RNA was isolated from plasma samples using the Maxwell 16 Viral Total Nucleic Acid Purification Kit (Promega). cDNA was synthesized using the SuperScript III First-Strand Synthesis System (Invitrogen) with a gag-specific primer: 5’-GCACTYAAGGCAAGCTTTATTGAGGCTTA-3’. The C2/V3 region (~416bp) of HIV *env* was amplified using a nested PCR approach with Phusion High-Fidelity PCR Master Mix (New England Biolabs). The outer primers were: 5’-ATTACAGTAGAAAAATTCCCCT-3’ and 5’-CAAAGGTATCCTTTGAGCCAAT-3’. The inner primers were: 5’-<u>TCGTCGGCAGCGTCAGATGTGTATAAGAGACAG</u>GAACAGGACCAGGATCCA ATGTCAGCACAGTACAAT-3’ and 5’-<u>GTCTCGTGGGCTCGGAGATGTGTATAAGAGACAG</u>GCGTTAAAGCTTCTGGG TCCCCTCCTGAG-3’, where the underlined portions indicate the Illumina adapter sequence. A unique barcode was added to each amplicon using the Nextera XT Index Kit (Illumina) and the barcoded amplicons were mixed to generate a sequencing library. Paired-end sequencing (2×300 bp) was performed using the Illumina MiSeq instrument and the MiSeq Reagent Kit v3.

### IGH amplification and ultra-deep sequencing

Total RNA was extracted from PBMCs using the Qiagen RNeasy Mini Kit. To reverse transcribe, amplify IGH encoding RNA, and generate sequencing-ready libraries, we used iRepertoire’s long read iR-Profile Kit and followed the procedure as described in the accompanying protocol (Wang et al., 2010). Paired-end sequencing (2×300 bp) was performed using the Illumina MiSeq instrument and the MiSeq Reagent Kit v3.

### HIV sequence data QC

Sequences from different samples were de-multiplexed by barcode using the internal software on the Illumina machine. We used the software package pRESTO (Vander Heiden et al., 2014) to assemble read pairs, remove sequences shorter than 300bp, remove sequences with a mean quality score less than 30, mask the primer sequences, and remove sequences that only occur once in a given sample. We then use an in house implementation of BLAST (Altschul, Gish, Miller, Myers, & Lipman, 1990) to check that each sequence has at least 70% identity to at least one HIV subtype *env* reference sequence, which were downloaded from LANL (Foley et al., 2017). In order to check for possible contaminations from HIV sequences outside of our study, we again used BLAST to map each of our sequences to every *env* sequence within the LANL database. In this case, any of our sequences that had 99% identity or more to any sequence within the *env* database would be deemed a contaminant. We found that all the samples from participant 8 in our study had a significant amount of identity with sequences derived from patient ID: 9036 in the LANL database. We also found that all the samples from our participant 9 had significant identity with sequences in the LANL database derived from patient ID: 9018. There are two reasonable explanations for this: i) these samples had a large degree of contamination, or ii) that our participants 8 and 9 are the same as patients 9036 and 9018 in the LANL database, respectively. We conclude that the latter is the more likely explanation because of the following rationale. This large degree of ‘contamination’ only occurred in participants 8 and 9, and it occurred in all their samples, however, the samples from these participants were processed in different batches. These participants’ diversity, and divergence trajectories showed a relatively steady increase over time, which would not be consistent with contamination (see results). Lastly, patients 9036 and 9018 from the LANL database both correspond to the study (Sturdevant et al., 2015) which also recruited participants from San Francisco, CA. None of the other samples in our study had detectable contamination using this method.

To check for cross contamination of sequences across samples in our study, we used a clustering approach. We first reduced the size of the dataset by grouping the sequences within each sample that have an edit distance less than or equal to 4 (see “Clustering sequences with samples” section below). We then choose the sequence within each group (or cluster) that had the highest count to be the ‘representative sequence’ for that cluster, after which we pooled all the representative sequences across all samples and clustered those pooled representative sequences using the same clustering algorithm. To identify clusters of sequences that were likely cross contaminants we used the following criteria: the cluster had to i) have a representative sequence that clustered closer with sequences from a different participant than with sequences from the same sample, and ii) have a frequency less than 0.001 within its sample. All sequences within clusters that satisfied these criteria were removed. This effectively identified low frequency sequences that were closer in genetic distance to sequences from another participant. Lastly, we used a phylogenetic approach (see “Making phylogenetic trees” section below) to i) make phylogenetic trees of the representative sequences in each sample, and ii) remove any sequence that is more closely related to a representative sequence that was identified as a cross contaminant (as described above) then to the other representative sequences in the sample (Figure S17).

### Clustering sequences within samples

We use the Needleman-Wunsch algorithm (Needleman & Wunsch, 1970) as implemented in the ‘needle’ program from the European Molecular Biology Open Software Suite (Rice, Longden, & Bleasby, 2000) to globally align each pair of sequences, and calculate the edit distances. Through an in-house algorithm, we then grouped sequences into a cluster that had an edit distance less than some provided threshold to any other sequence in the cluster. For HIV sequences, this threshold was 4; for AbR sequences, this threshold was 6.

### Phylogenetic analysis

To make a phylogenetic tree of a group of sequences, we first made multiple sequence alignments using MAFFT (Katoh et al., 2002), and then constructed phylogenetic trees using FastTree (Price, Dehal, & Arkin, 2010). Visualization and analyses of newick formatted files was performed using the ETE toolkit (Huerta-Cepas, Serra, & Bork, 2016).

### Calculating HIV divergence

We first assigned an HIV reference sequence for each participant by finding the most abundant sequence at the first time-point. Because participant 6 showed extensive evidence of a super-infection occurring at the second time-point, we assigned two reference sequences to participant 6: the most abundant sequences from the first and second time-points. We then translated the reference sequences as well as all other sequences in the data (see “Translating HIV sequences” section below). To find the number of synonymous and non-synonymous changes for a given query HIV sequence, we first codon aligned it to the participant’s reference. For a given codon, we first calculated the number of expected non-synonymous and synonymous sites as:

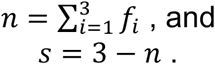

Where *f*_*i*_ gives the proportion of all possible nucleotide changes at codon position *i* of the reference sequence that result in an amino acid change. We denote *N* and *S* as the sum of *n* and *s* across all codons in a given reference sequence. We then counted the number of observed non-synonymous and synonymous changes in a given query HIV sequence as *N*_*o*_ and *S*_*o*_. If there were multiple mutations, we selected the order of mutations that resulted in the least amount of amino acid changes as the most parsimonious, and thus most likely. The proportion of non-synonymous and synonymous mutations in a given query sequence is then,

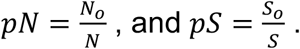

To estimate non-synonymous and synonymous divergence we then use

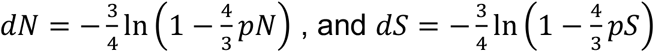

(Nei & Gojobori, 1986). The notation above was heavily borrowed from Richard Orton’s blog post (Orton, 2014). Because participant 6 had two reference sequences, divergence for a given query sequence was calculated relative to both reference sequences, and the lower value was used.

### Calculating HIV selection

Selection in a given HIV sequence was estimated by taking the ratio of the non-synonymous over synonymous divergence values (described above), *dN*/*dS*.

### Translating HIV sequences

In order to translate a given query HIV sequence, we first used needle to globally align it to the reference HXB2 *env* sequence (downloaded from LANL). We then used this alignment to determine the coding frame of the query sequence. Once this was known we translated the query nucleotide sequence using a simple in-house python script.

### Calculating diversity

To estimate diversity in both the HIV population and the AbR we calculated the statistic, *π* (Strauli & Hernandez, 2016). In words, *π* is the expected genetic distance between two randomly selected sequences from a given sample. Mathematically *π* can be expressed as

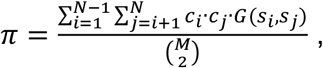

where *N* gives the total number of unique sequences in the sample, *c*_*i*_ gives the count of sequence *i*, *M* gives the total counts of sequences in that sample (i.e. 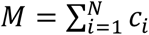), and *G*(*x, y*) gives the genetic distance between sequences *x* and *y*. We used VSEARCH with the “--allpairs_global” option to globally align all pairs of sequences in a sample (Rognes, Flouri, Nichols, Quince, & Mahé, 2016). Genetic distance between a pair of sequences was then calculated as the percent of mismatches in the alignment.

### AbR sequence data QC

As with the HIV data, we used pRESTO to assemble read pairs, remove reads less than 300bp, remove reads with a mean Q score less than 20, and remove reads that only occur once. We then use IgBLAST (Ye, Ma, Madden, & Ostell, 2013) to align each sequence to a database of germline immunoglobin genes downloaded from the IMGT website (imgt.org) (Giudicelli et al., 2006). After this, we used Change-O to annotate each sequence with its most likely V, D, and J germline gene-segments, identify the FWRs and CDRs, and to construct the likely naïve antibody sequence from the germline gene-segment alignments.

### Calculating Ab divergence

Divergence in a given Ab sequence was calculated as the number of changes in the observed sequence relative to the naïve sequence, divided by the length of the sequence. This is excluding the junction region of the sequence, as naïve sequence reconstruction of this region is unreliable.

### Estimating Ab Selection

We ran the BASELINe program on each individual Ab sequence and used the resulting ∑ value to estimate selection pressure. For a detailed description of this tool see (Yaari et al., 2012). Very briefly, BASELINe compares the observed number of mutations in a sequence (relative to its inferred naïve ancestor) to a null distribution of the expected number of mutations under no selection. The program takes local nucleotide motifs into account when calculating mutation probabilities and returns a ∑ value that indicates the distance between the observed number of mutations and the null distribution. A negative ∑ indicates fewer mutations than expected (negative selection), and a positive ∑ indicates more mutations than expected (positive selection). It does this separately for different regions of the Ab sequence (i.e. the FWR and CDR).

### Creating AbR lineages

To cluster the AbR of a given participant, we first divided it into partitions by grouping together all sequences that use the same germline V and J gene-segments (as annotated by Change-O). We then clustered the sequences within a given partition/time-point (see “Clustering sequences within samples” section above), with an edit distance threshold of 6 (Figure S18 A). Once clusters were delineated, and similar to the clustering of HIV sequences, we assigned the most numerous sequence of each cluster as the ‘representative sequence’. We then linked clusters, within a given partition, across adjacent time-points using the following algorithm: We first found the representative sequence in the previous time-point that had the smallest edit distance to a given representative sequence in a contemporary time-point. If this edit distance was smaller than 30 (Figure S18 B), then the two representative sequences (and the clusters they represented) were linked as being part of the same lineage. This process was carried out independently in each participant, over each representative sequence, and for each adjacent time-point pair (see Figure S19 A). Finally, once all lineages were assigned in this manner, any lineage that did not rise above 0.0001 frequency, in any of the time-points was disregarded.

### Simulating null AbR lineages

To simulate random AbR lineages, we began by using our clustered Ab sequences within the partitioned AbR data (see “Creating AbR lineages” section above). We then carried out an identical procedure as was done to create the observed lineages, with the exception that instead of finding the parent cluster in the previous time-point that had the minimum edit distance, a parent cluster was randomly chosen from the previous time-point (Figure S19 B). Additionally, in order to replicate the aspect of the observed data where each time-point brought a certain number of new lineages to the population, we estimated the probability of a new lineage as

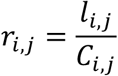

where *l*_*i,j*_ gives the number of new lineages, and *C*_*i,j*_ gives the total number of clusters in participant, *i*, and time-point, *j*. For the simulated lineages, we then randomly assigned each cluster as being a new lineage (i.e. not having any connections with the previous time-point) with probability *r*_*i,j*_. In order to have a null dataset that was sufficiently large, we duplicated the observed, clustered data N times and then simulated lineages using this N-fold larger dataset. When comparing all lineages to simulated data (i.e. Figure 8), N=10; when comparing lineages from particular AbR partitions to simulated data (i.e. Figure 7), N=100.

### Linear modeling of population level interactions

We tested for cross-participant, population-wide interactions of summary statistics using a linear mixed model approach. We modeled the interaction of a pair of summary statistics as

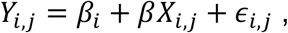

where *Y*_*i,j*_ gives the value of a given HIV summary statistic for the *j*th time-point of participant *i*, *X*_*i,j*_ gives the value of a given AbR summary statistic for the same participant/time-point, *β*_*i*_ is a random intercept term to correct for participant specific effects, and *∈*_*i,j*_ is a random error term that is assumed to be normally distributed with a mean of 0. This model was implemented in the R programming language using the ‘lmer’ function of the ‘lme4’ package (Bates, Mächler, Bolker, & Walker, 2015). We then used a likelihood ratio test to determine if a model with *β*≠0 provided a significantly better fit than a model with *β*=0 (p≤0.05). If it did, then the given pair of HIV and AbR summary statistics was deemed to be interacting.

### Trajectory permutation test

We test for associations between a set of AbR trajectories to one or more HIV trajectories using a permutation-based test. The AbR and HIV trajectories were loaded into memory as matrices where each row is a different trajectory and each column is a time-point in chronological order from left to right. We first standardize each trajectory by subtracting the mean and dividing by the standard deviation. If 𝑣 is a row vector representing a given trajectory, then 𝑣 is standardized by

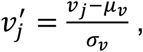

where *μ_𝑣_* and *σ_𝑣_* give the arithmetic mean and standard deviation of 𝑣, respectively. We then calculate the sum of the squared error (SSE) for a given AbR trajectory relative to a given HIV trajectory as

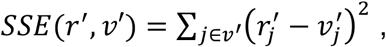

where *r*′ is a standardized HIV trajectory vector. This gives the observed SSE values for each AbR/HIV trajectory pair. If either the HIV or AbR trajectories have missing values, then these time-points are disregarded in the SSE calculation, and a trajectory must have at least 75% of its values defined to be included in the test. We then permute the columns of the AbR trajectory matrix many times and calculate the SSE values for each permuted AbR trajectory after each permutation. This gives our permuted null distribution of SSE values. If an observed AbR/HIV trajectory pair have an SSE value that is significantly outside of this null distribution, then they are deemed to be significantly associated with one another, where significance is appropriately adjusted as based on the number of tests. When conducting this test on whole AbR population trajectories vs. whole HIV population trajectories (Figure 4 B-E), we performed 100,000 permutations for each participant. When conducting this test on AbR partition trajectories vs. whole HIV population trajectories (Figures 5, S12, and S13) we performed 1,000 permutations for each participant.

### Comparison against published datasets

To compare the results of our trajectory permutation test to a literature dataset we used a Mann-Whitney U test. However, before this could be done, we first must combined the results of our permutation-based test across participants. When the permutation-based test was employed to identify HIV associated AbR partitions, the structure of the results was as follows: each participant had hundreds of AbR partitions, and each AbR partition had tens of p values associated with it (from comparing each of its summary statistics to each of the HIV population summary statistics). In order to combine p values such that there was one value associated with one V gene segment, we first pooled the p values across all AbR partitions that have a given V gene segment, and across all participants. We then used Fisher’s method to arrive at an overall V gene score for this pool of p values. If *p* is a vector of p values associated with a given V gene segment, then we first combine the p values into one overall p value, *p*_*overall*_, using Fisher’s method:

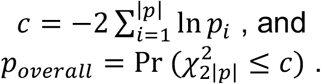

Where 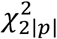 is the chi-squared distribution with 2|*p*| degrees of freedom. Strictly speaking, *p*_*overall*_ will tend to be inflated because not all the p values in *p* are independent (ex. AbR partition trajectories that are associated with HIV selection, will also tend to be associated HIV non-synonymous divergence). However, because this inflation should be the same across V gene segments, and because we were not interested in actual significance but rather needed some reasonable method for combining p values into an overall score, Fisher’s method should be sufficient. Finally, to avoid *p*_*overall*_ being interpreted as a significance level, we took its log transform to arrive at a V gene score

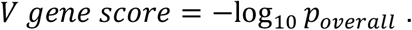

We then used a Mann-Whitney U test to see if V gene segments that were ‘well represented’ in a literature dataset tended to have significantly different *V gene score* values then those that were not ‘well represented’. In the case of the dataset of HIV targeting Abs, ‘well representation’ was defined as presence/absence (i.e. count ≥ 1). The dataset of influenza targeting Abs was relatively large (432 entries), so ‘well representation’ was defined as a count ≥ 10 (Table S1).

### Calculating mutual information (MI)

To measure the amount of association between two sites (columns) in a pair of MSAs we first reduced the complexity of the amino acid code by converting it to a code of ‘change’ or ‘no-change’. In this case, if a site had an amino acid identity that was different than the previous time-point, then it was recorded as a ‘1’ and if it was the same, then it was recorded as a ‘0’ (the first time-point is always ‘0’). We then used MI to measure the amount of association (coevolution) between two columns in a ‘change’, ‘no-change’ alignment. MI is calculated as

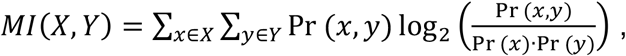

where *X* and *Y* are categorical random variables representing the two different columns being compared. *x* and *y* represent particular states of *X* and *Y*, respectively (i.e. ‘1’ or ‘0’ for a change/no-change alignment). Pr(*a*) is the probability that a given random variable (or MSA column) equals the state, *a*. If *c* is a vector that represents a given column of an MSA, then Pr(*a*)can be estimated as

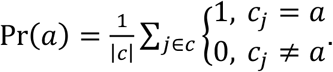

Pr(*a, b*) is the joint probability that the random variable representing one MSA column equals *a*, and simultaneously the random variable representing the other MSA column equals *b*. If *d* is a vector that represents a given column of the other MSA, then Pr(*a, b*) can be estimated as

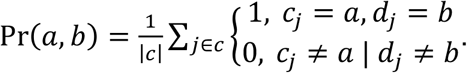

### Correcting for multiple tests

Unless otherwise stated, p values from a given statistical test within a participant were corrected for multiple testing using the Benjamini-Hochberg procedure.

### Network plots

Networks of lineages were visualized using cytoscape (Shannon et al., 2003).

### Software

All computer code written for this study is available on GitHub, at https://github.com/nbstrauli/abr_hiv_evo. This is accompanied by a flow chart which shows each data-processing step with the corresponding name of the script that carries it out (Figure S20).

### Data availability

All sequence data associated with this study is available at the sequence read archive (SRA) under the BioProject ID: PRJNA543982.

### Website

To create an interactive web application, we built a website using the Boostrap framework for the frontend interface along with the JavaScript graphing library plotly.js. The website is available at https://ab-hiv-coevolution.github.io/HIV_AB_CoEvo/

## Acknowledgements

First and foremost, we would like to thank the anonymous participants who donated their data, time, and person to make this study (and those like it) possible. We will never know who you are, but we are eternally grateful for your contributions. We thank Dr. Manon Eckhardt for her edits and figure contributions. We would like to thank Alf Seccombe for driving a poor graduate student across town in order to get MiSeq kits into a proper freezer. Lastly, We thank Dr. Kenneth Hoehn for valuable manuscript feedback. This research was supported by grants from the National Institutes of Health (R01HG007644 to RDH), and University of California, San Francisco-Gladstone Institute of Virology & Immunology Center for AIDS Research (P30-AI027763).

