## Supplemental Figures for "The genetic interaction between HIV and the antibody repertoire" (from another version of this preprint: version 2, 2019-05-24)

### Title

**Supplementary Figures**

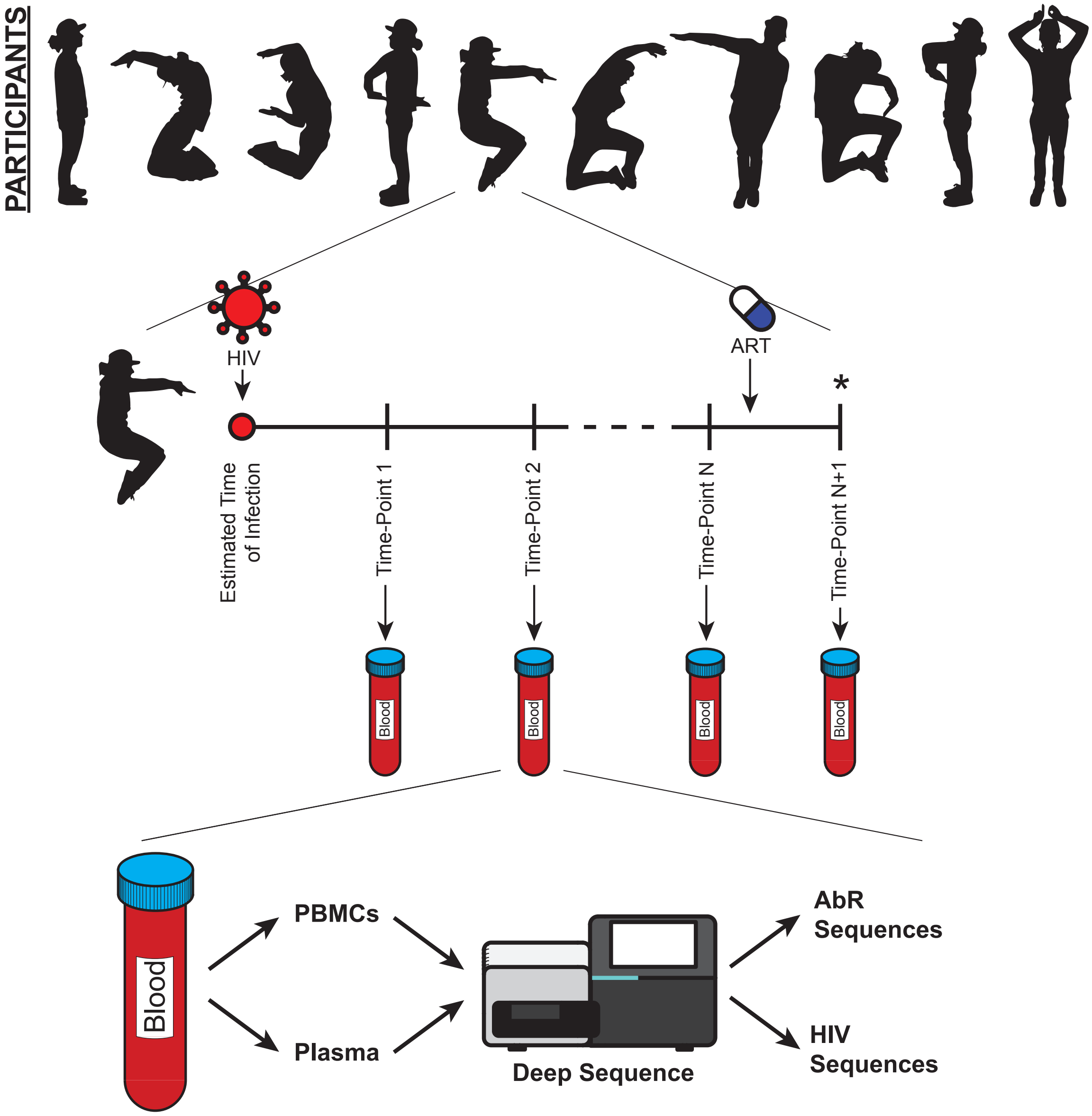


**Figure S1. Schematic of study.**
There were 10 participants in our study. Peripheral blood was drawn from each participant, post HIV infection, for 10-20 longitudinal time-points. Each blood sample was divided into PBMCs and plasma. Ab sequences were derived from the PBMCs and HIV sequences were derived from plasma. Note that participant avatars in no way reflect actual participants and instead reflect one lonely scientist.
*Only participants 1, 2, and 5 had the final time-point with ART exposure.

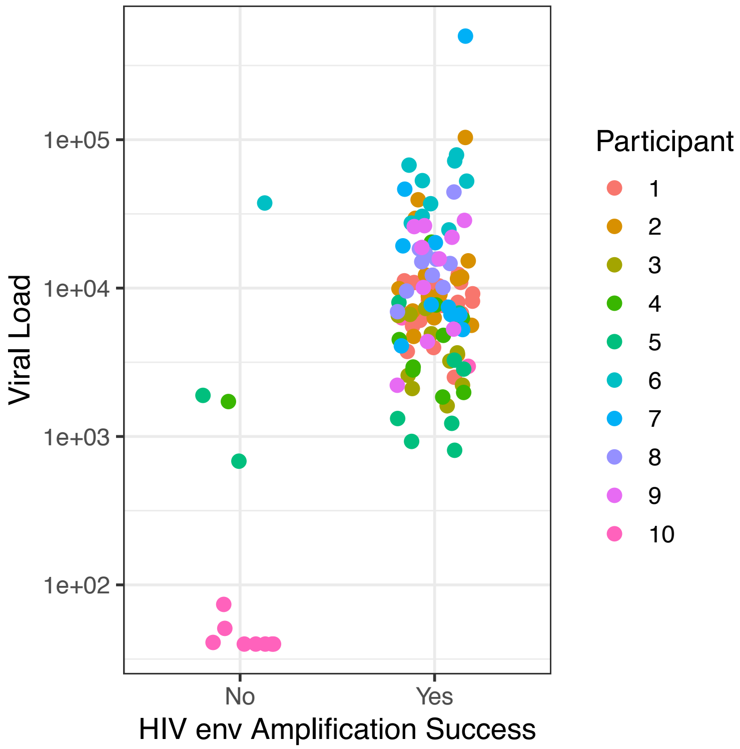


**Figure S2. HIV amplification success and viral load.**
Depicts the relationship between HIV viral load and our ability to amplify HIV *env* prior to sequencing. All samples in which *env* could not be amplified had low to exceptionally low viral load, with the exception of one sample from participant 6.

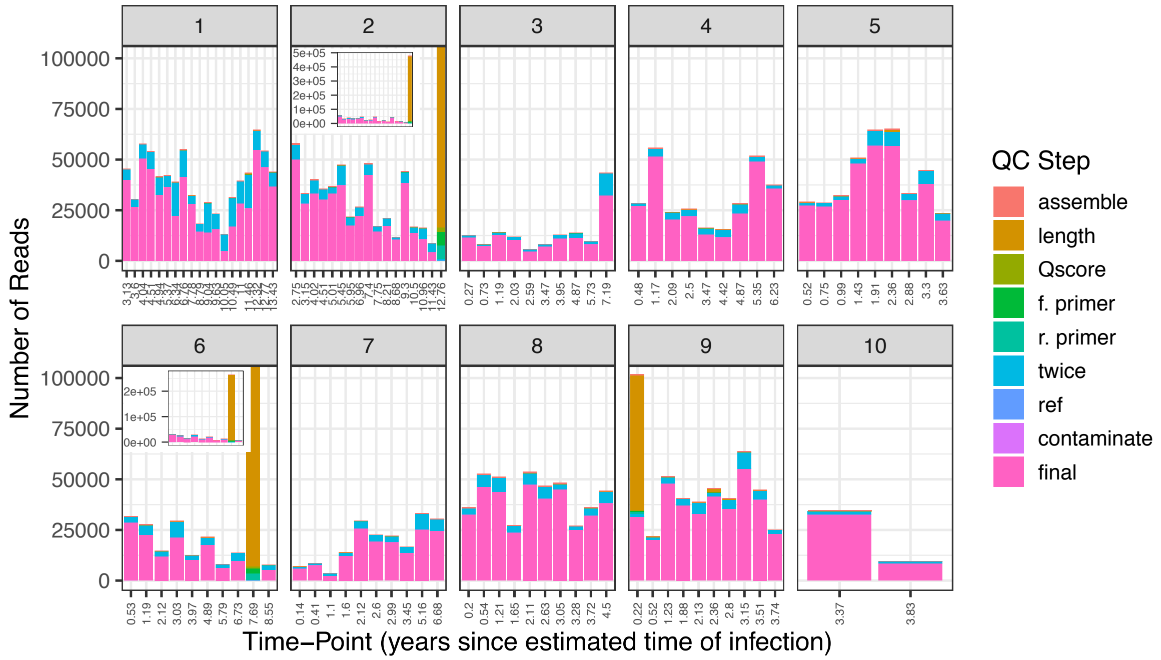


**Figure S3. Read depth and QC filtering for HIV data.**
Stacked bar charts depicting the total read depth and proportion of reads that were filtered after each QC step. The height of each bar shows initial read depth (directly after sequencing) for each HIV sample. The relative lengths of each color within a bar show how many reads were filtered out from a given QC step. The legend maps color to QC step, and QC steps are shown (from top to bottom) in the order that they were performed: “assemble” = could not assemble read pairs; “length” = too short; “Qscore” = mean Q score too low; “f. primer” = could not align forward primer; “r. primer” = could not align reverse primer; “twice” = sequence did not occur at least twice; “ref” = did not align to an HIV reference sequence; “contaminate” = sequence was found in other studies; “final” = the final sequences that passed all QC. Numeric titles for each panel give the participant ID. Insets in participants 2 and 6 show the full breadth of the y-axes, depicting the data from two samples that did not successfully amplify *env*, but were included as a negative control. They did not pass QC.

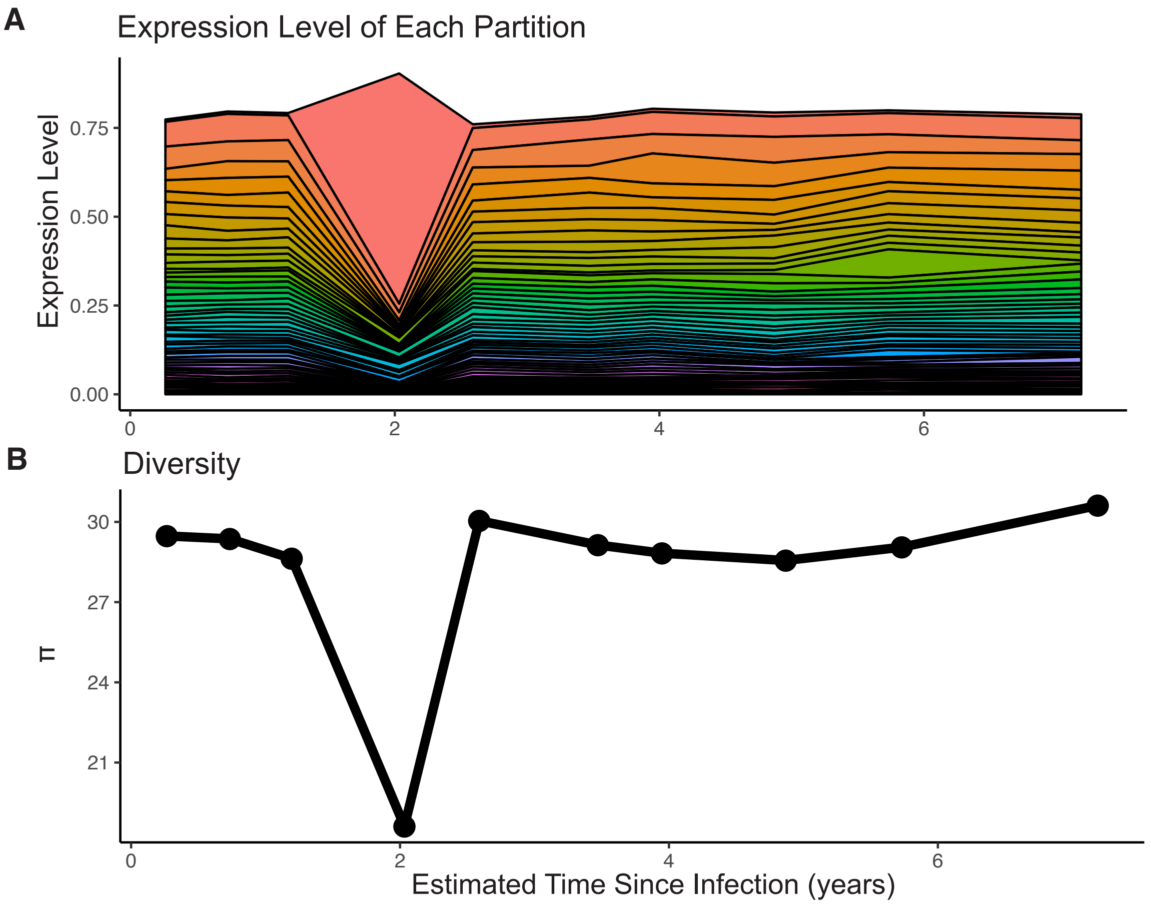


**Figure S4. Anomalous time-point in participant 3.**
(A) Stacked area chart shows the unrealistic clonal expansion occurring at the 4$^{th}$ time-point of participant 3. Each unique color represents a unique AbR partition, and the width of a given color represents the relative frequency of that AbR partition at that time-point. (B) A line plot showing the corresponding dramatic drop in AbR diversity due to this anomaly.

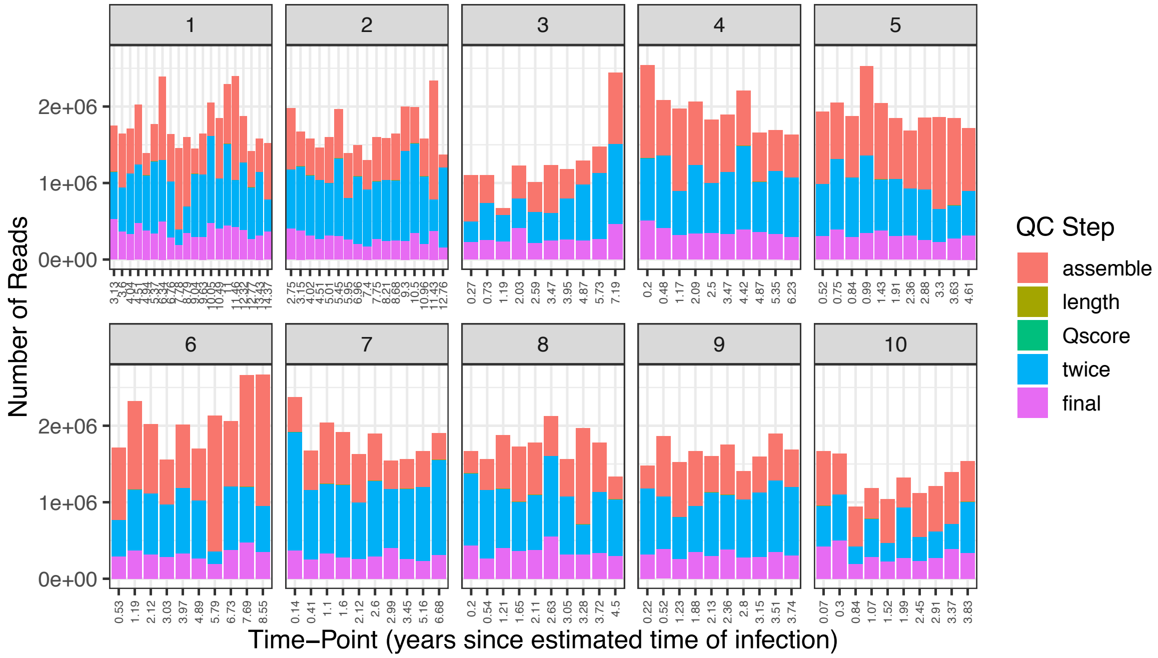


**Figure S5. Read depth and QC filtering for AbR data.**
Stacked bar charts depicting the total read depth and proportion of reads that were filtered after each QC step. The height of each bar shows initial read depth (directly after sequencing) for each AbR sample. The relative lengths of each color within a bar show how many reads were filtered out from a given QC step. The legend maps a color to a QC step, and QC steps are shown (from top to bottom) in the order that they were performed: “assemble” = could not assemble read pairs; “length” = too short; “Qscore” = mean Q score too low; “twice” = sequence did not occur at least twice; “final” = the final sequences that passed all QC. Numeric titles for each panel give the participant ID.

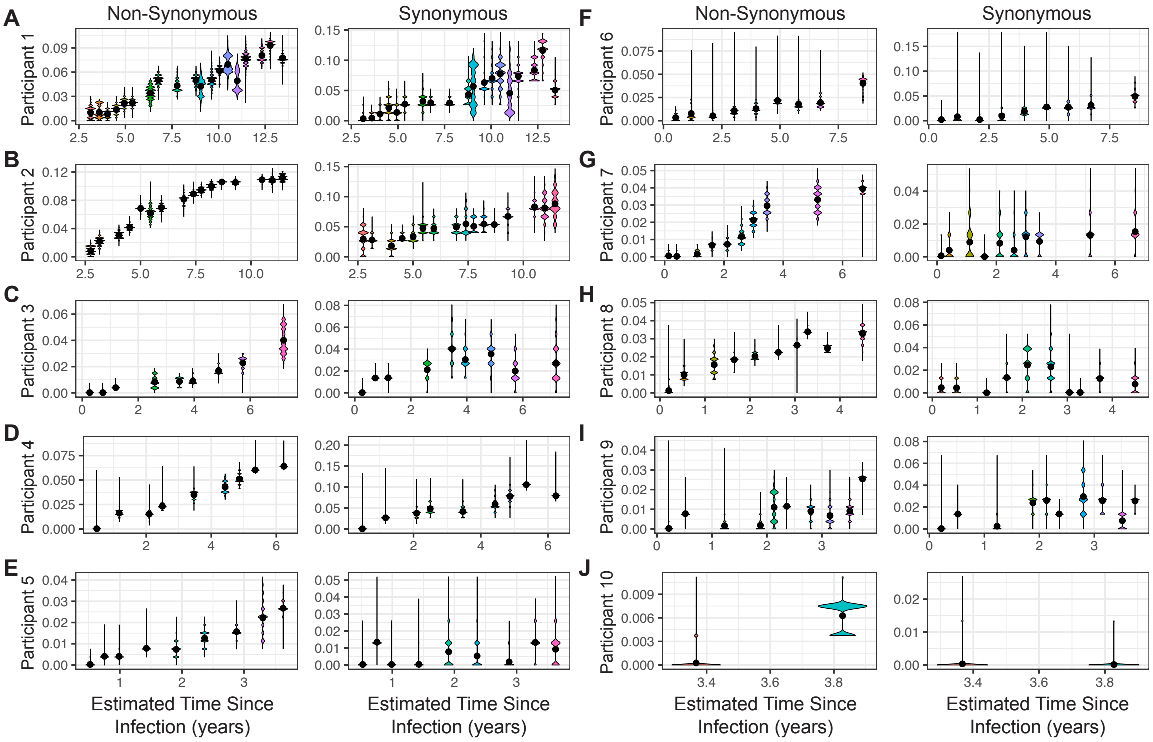


**Figure S6. Distributions of HIV divergence values.**

Each violin plot gives the distribution of HIV divergence values for a given time-point, where black points show the mean of the distribution. Plots on the left side of a panel show non-synonymous divergence, and plots on the right show synonymous divergence. (A) Participant 1. (B) Participant 2. (C) Participant 3. (D) Participant 4. (E) Participant 5. (F) Participant 6. (G) Participant 7. (H) Participant 8. (I) Participant 9. (J) Participant 10.

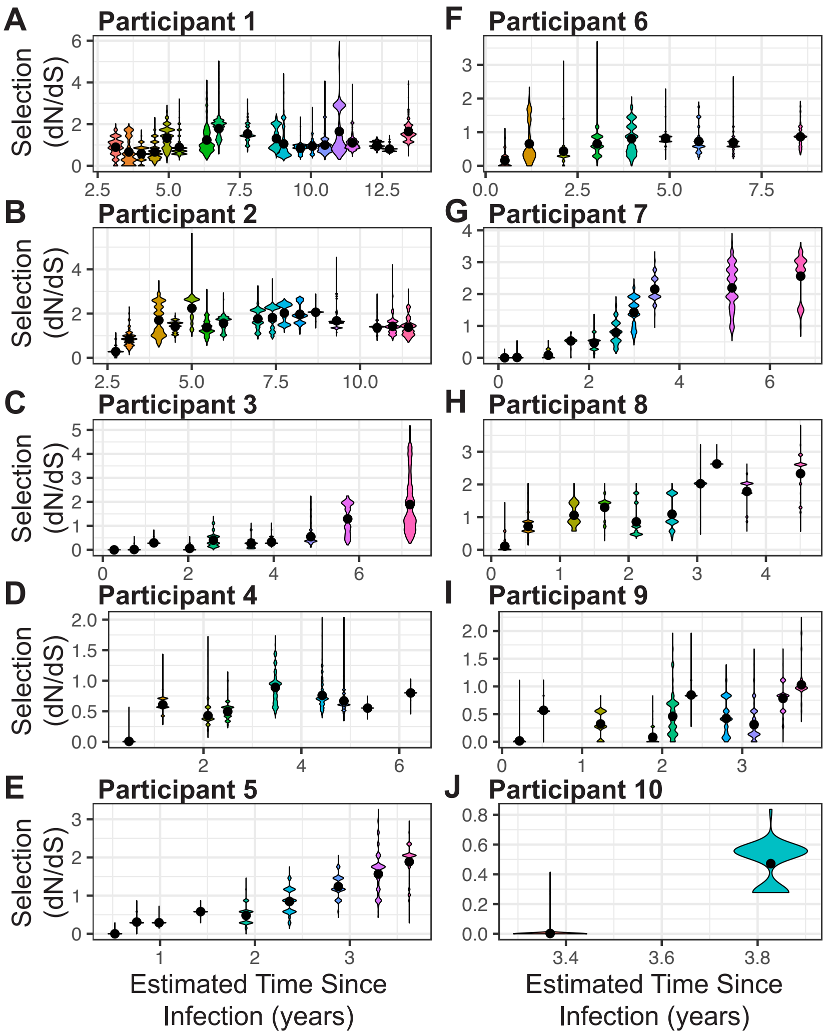


**Figure S7. Distributions of HIV selection values.**

Each violin plot gives the distribution of HIV selection values at a given time-point, where black points show the mean of the distribution. (A) Participant 1. (B) Participant 2. (C) Participant 3. (D) Participant 4. (E) Participant 5. (F) Participant 6. (G) Participant 7. (H) Participant 8. (I) Participant 9. (J) Participant 10.

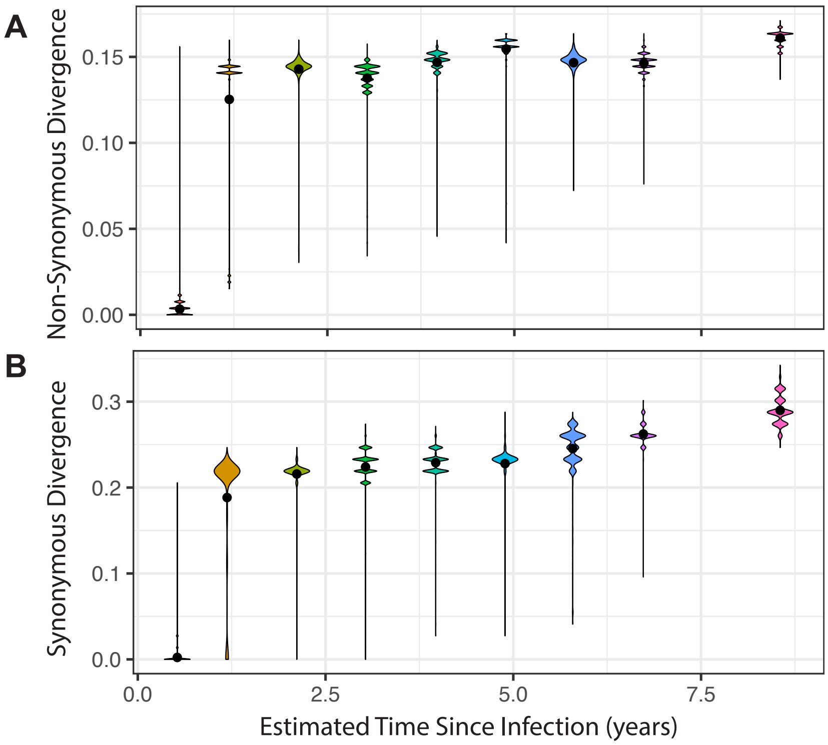


**Figure S8. Super infection in participant 6.**

Each violin plot gives the distribution of HIV divergence values for a given time-point for participant 6, where black points show the mean of the distributions. Divergence was calculated here in the same way as other participants in order to illustrate super-infection. (A) Non-synonymous divergence. (B) Synonymous divergence. The large increases in both synonymous and non-synonymous divergence between the 1st and 2nd time-points are indicative of a super-infection.

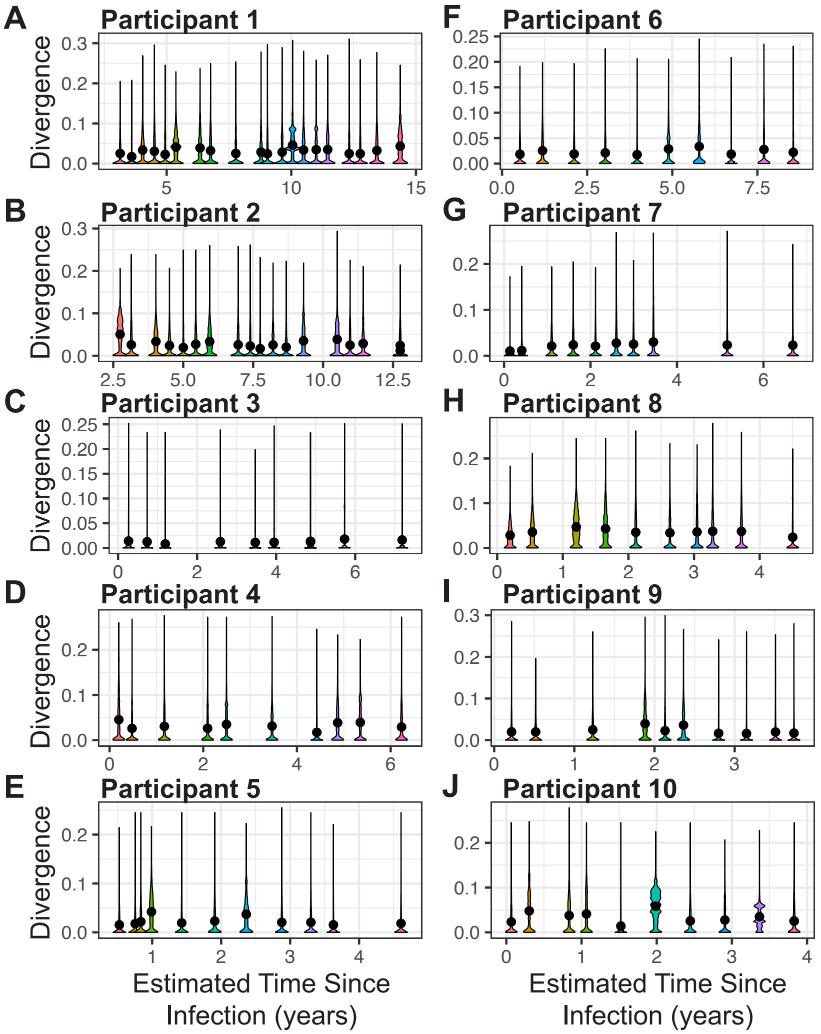


**Figure S9. Distributions of antibody divergence values.**
Each violin plot gives the distribution of antibody divergence values for a given time-point, where black points show the mean of the distribution. (A) Participant 1. (B) Participant 2. (C) Participant 3. (D) Participant 4. (E) Participant 5. (F) Participant 6. (G) Participant 7. (H) Participant 8. (I) Participant 9. (J) Participant 10.

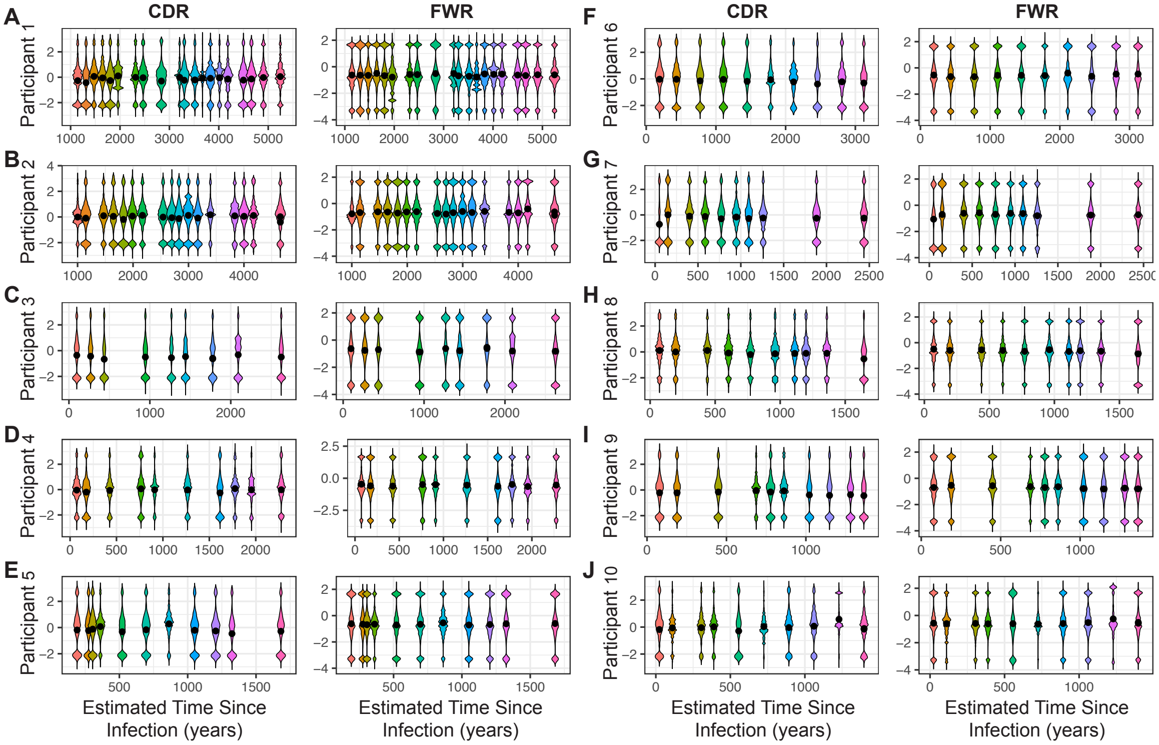


**Figure S10. Distributions of AbR selection values.**

Each violin plot gives the distribution of AbR selection values at a given time-point, where black points show the mean of the distribution. Plots on the left side of a panel show selection in the CDR, and plots on the right show selection in the FWR. (A) Participant 1. (B) Participant 2. (C) Participant 3. (D) Participant 4. (E) Participant 5. (F) Participant 6. (G) Participant 7. (H) Participant 8. (I) Participant 9. (J) Participant 10.

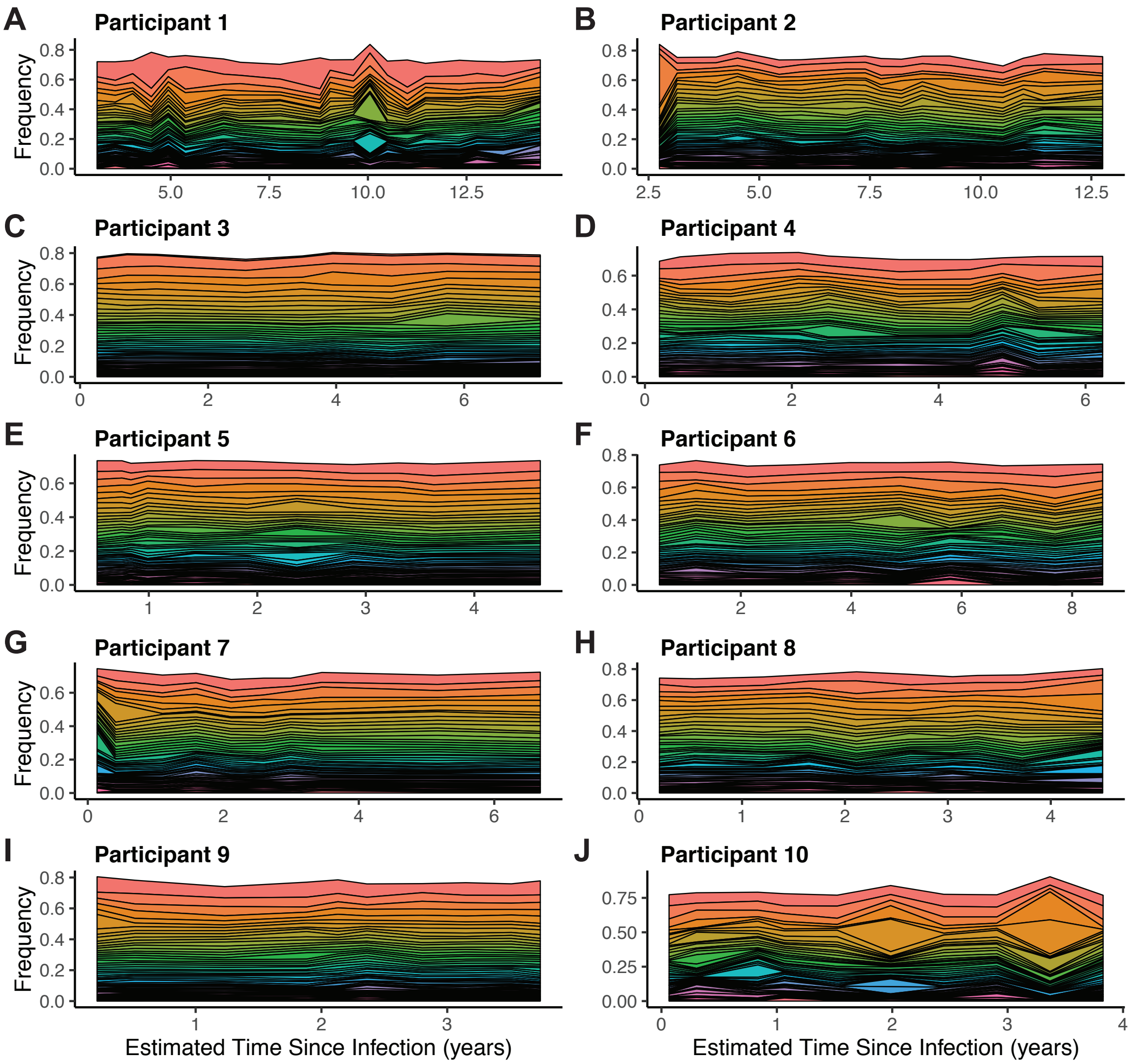


**Figure S11. AbR partition frequencies over time.**
Stacked area charts of AbR partition frequencies over time. Each unique color represents a unique AbR partition, and the width of a given color represents the relative frequency of that AbR partition at that time-point. Only the top 50 AbR partitions are shown, when sorted by the sum of frequencies across all time-points. (A) Participant 1. (B) Participant 2. (C) Participant 3. (D) Participant 4. (E) Participant 5. (F) Participant 6. (G) Participant 7. (H) Participant 8. (I) Participant 9. (J) Participant 10.

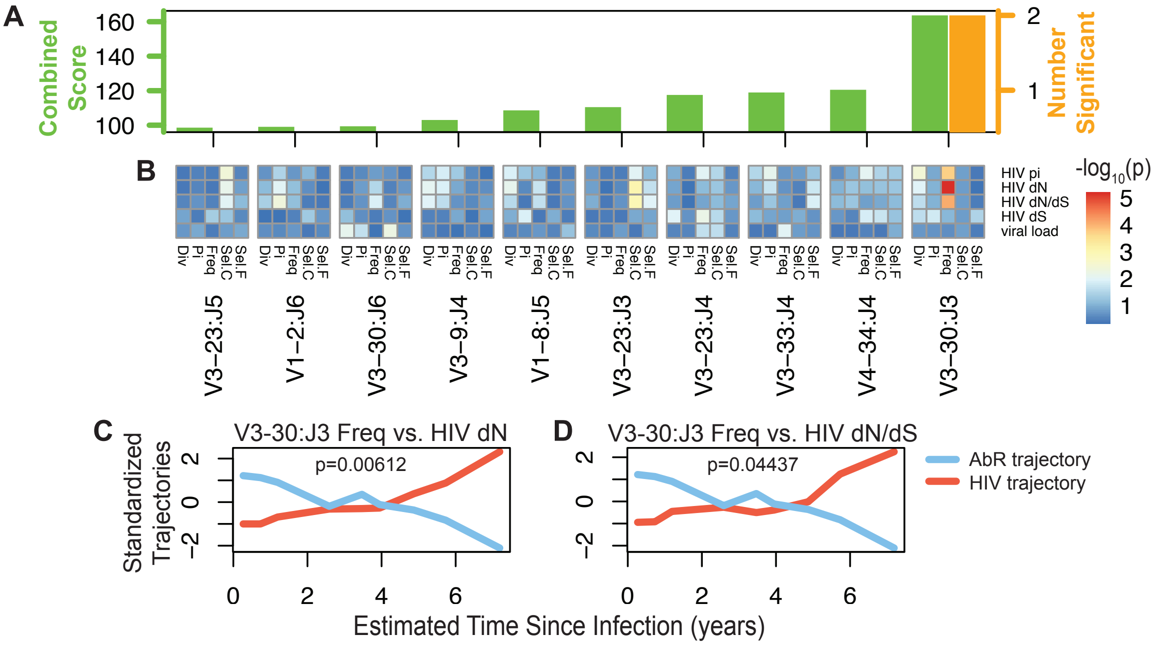


**Figure S12. Results of permutation-based test to identify HIV-associated AbR partitions in participant 3.**

(A) A barplot showing the combined score from the permutation-based test (left axis), and the number of significant associations (right axis) for the top 10 AbR partitions. AbR partitions were sorted first by the number of significant associations, then by their combined score from the permutation-based test (ascending, left-right). (B) Heatmaps depicting the significance ($-\log_{10}$(p-value)) foreach test run within each of the top 10 AbR partitions. Columns correspond to summary statistics of the AbR partitions: Div=divergence, Pi=diversity, Freq=relative frequency, Sel.C=CDR selection, and Sel.F=FWR selection. Rows correspond to summary statistics of the HIV population: HIV pi=diversity, HIV dN=nonsynonymous divergence, HIV dS=synonymous divergence, HIV dN/dS=selection, and viral load is self-explanatory. The color of each element in the heatmaps shows the significance of the association between a given AbR partition summary statistic trajectory with a given HIV population summary statistic trajectory. (C, D) Shows the AbR partition (blue) and HIV population (red) trajectories that were significantly associated. (C) Frequency trajectory of the IGHV3-30:IGHJ3 AbR partition with the divergence trajectory of the HIV population. (D) Frequency trajectory of the IGHV3-30:IGHJ3 AbR partition with the selection trajectory of the HIV population.

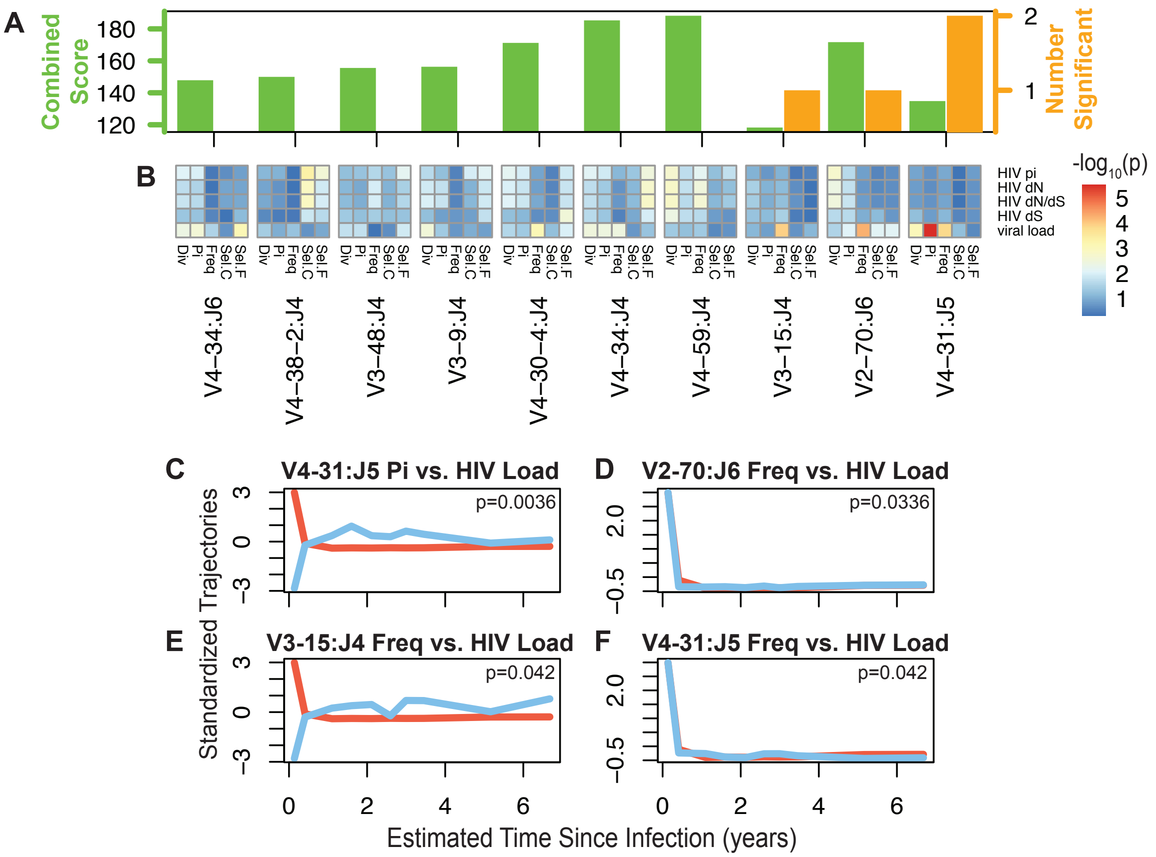


**Figure S13. Results of permutation-based test to identify HIV-associated AbR partitions in participant 7.**

(A) A barplot showing the combined score from the permutation-based test (left axis), and the number of significant associations (right axis) for the top 10 AbR partitions. AbR partitions were sorted first by the number of significant associations, then by their combined score from the permutation-based test (ascending, left-right). (B) Heatmaps depicting the significance (-$\log_{10}$(p-value)) foreach test run within each of the top 10 AbR partitions. Columns correspond to summary statistics of the AbR partitions: Div=divergence, Pi=diversity, Freq=relative frequency, Sel.C=CDR selection, and Sel.F=FWR selection. Rows correspond to summary statistics of the HIV population: HIV pi=diversity, HIV dN=nonsynonymous divergence, HIV dS=synonymous divergence, HIV dN/dS=selection, and viral load is self-explanatory. The color of each element in the heatmaps shows the significance of the association between a given AbR partition summary statistic trajectory with a given HIV population summary statistic trajectory. (C-F) Shows the AbR partition (blue) and HIV population (red) trajectories that were significantly associated. (C) Diversity trajectory of the IGHV4-31:IGHJ5 AbR partition with the viral load trajectory of the HIV population. (D) Frequency trajectory of the IGHV2-70:IGHJ6 AbR partition with the viral load trajectory of the HIV population. (E) Frequency trajectory of the IGHV3-15:IGHJ4 AbR partition with the viral load trajectory of the HIV population. (F) Frequency trajectory of the IGHV4-31:IGHJ5 AbR partition with the viral load trajectory of the HIV population.

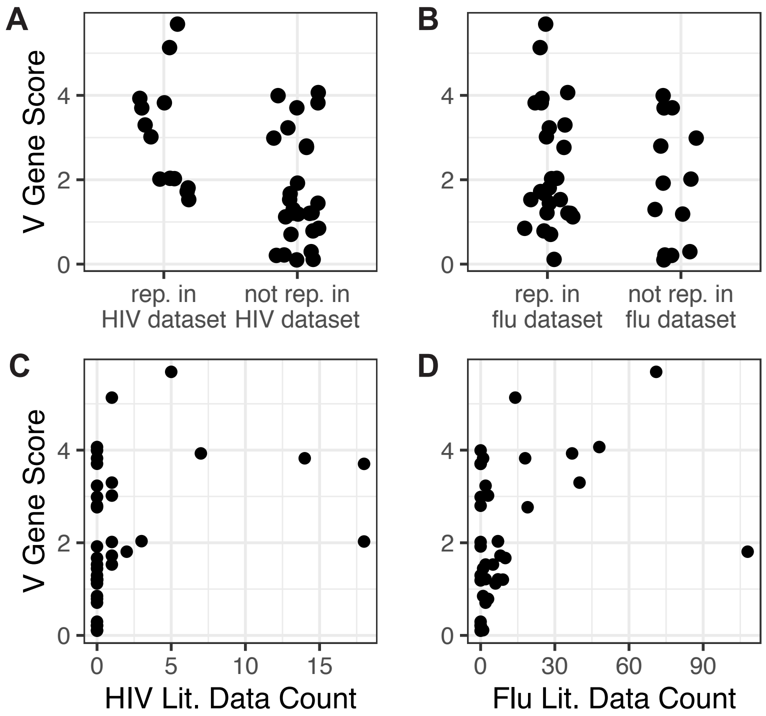


**Figure S14. Literature validation of HIV-associated AbR partitions.**
(A, B) Shows the overall score from our permutation-based test for each V gene segment when partitioned as being represented in a literature dataset (points to the left), vs. not represented in a literature dataset (points to the right). (A) Pertains to the HIV literature dataset, where 'representation' is defined as the V gene segment occurring at least once. (B) Pertains to the influenza literature dataset, where 'representation' is defined as the V gene segment occurring at least 10 times. (C, D) Scatter plots showing the relationship between the overall score from our permutation-based test for each V gene segment (y-axis) and the count of a given V gene segment in a literature dataset (x-axis). (C) HIV literature dataset. (D) Influenza literature dataset.

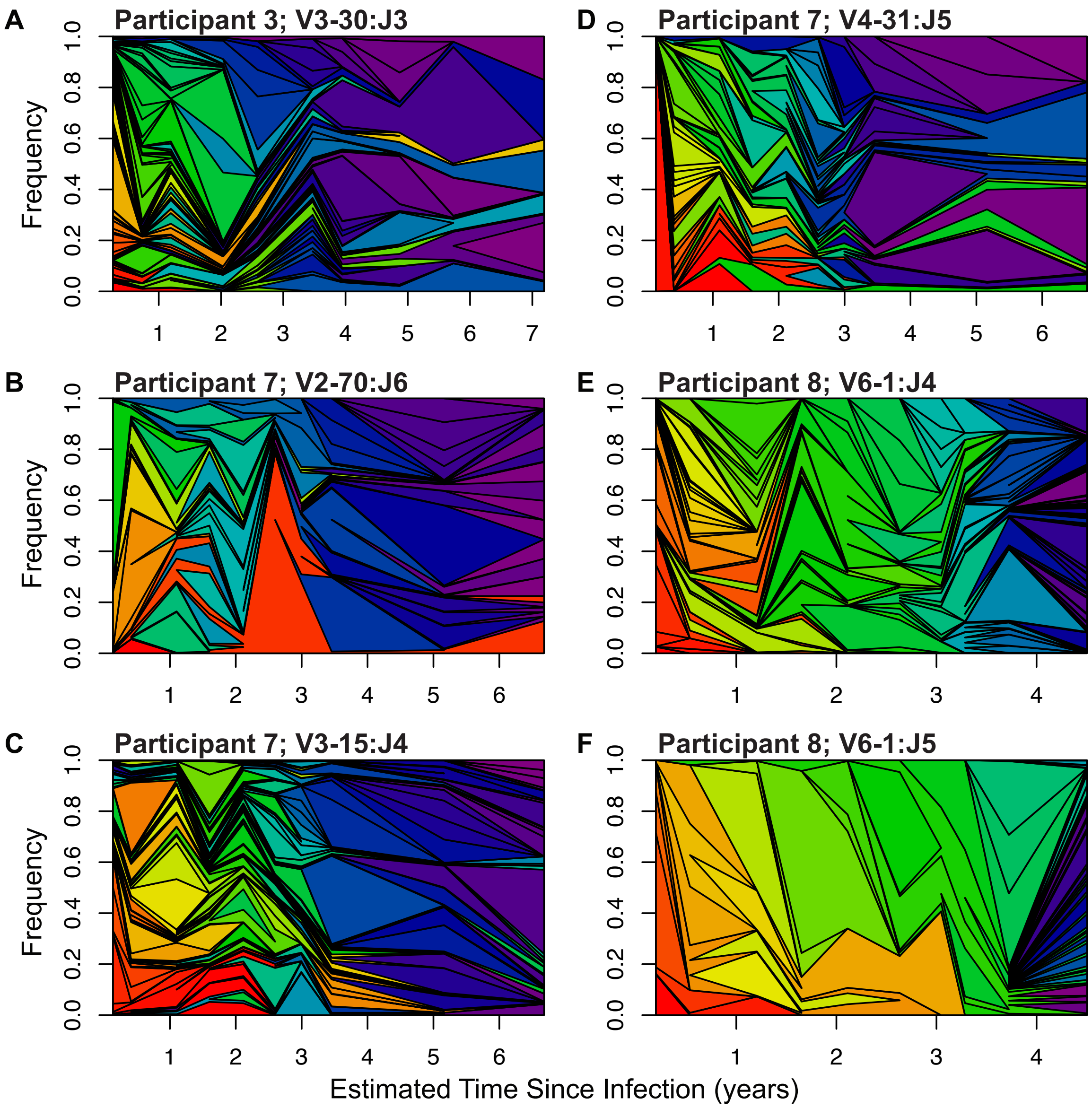


**Figure S15. Muller plots of Ab lineages.**

Depicts the relative frequency of each lineage within a given HIV-associated AbR partition. Each unique color represents a unique lineage. If a new lineage arises within the bounds of a preexisting lineage, then the new lineage is a daughter of the preexisting, parent lineage. Lineages that began earlier in the time-course have colors closer to the red side of the spectrum, while lineages that began later in the time-course have colors closer to violet. Only lineages that exceeded 0.0001 frequency in the larger AbR for at least one time-point are included in the plot. (A) AbR partition IGHV3-30:IGHJ3 in participant 3. (B) AbR partition IGHV2-70:IGHJ6 in participant 7. (C) AbR partition IGHV3-15:IGHJ4 in participant 7. (D) AbR partition IGHV4-31:IGHJ5 in participant 7. (E) AbR partition IGHV6-1:IGHJ4 in participant 8. (F) AbR partition IGHV6-1:IGHJ5 in participant 8.

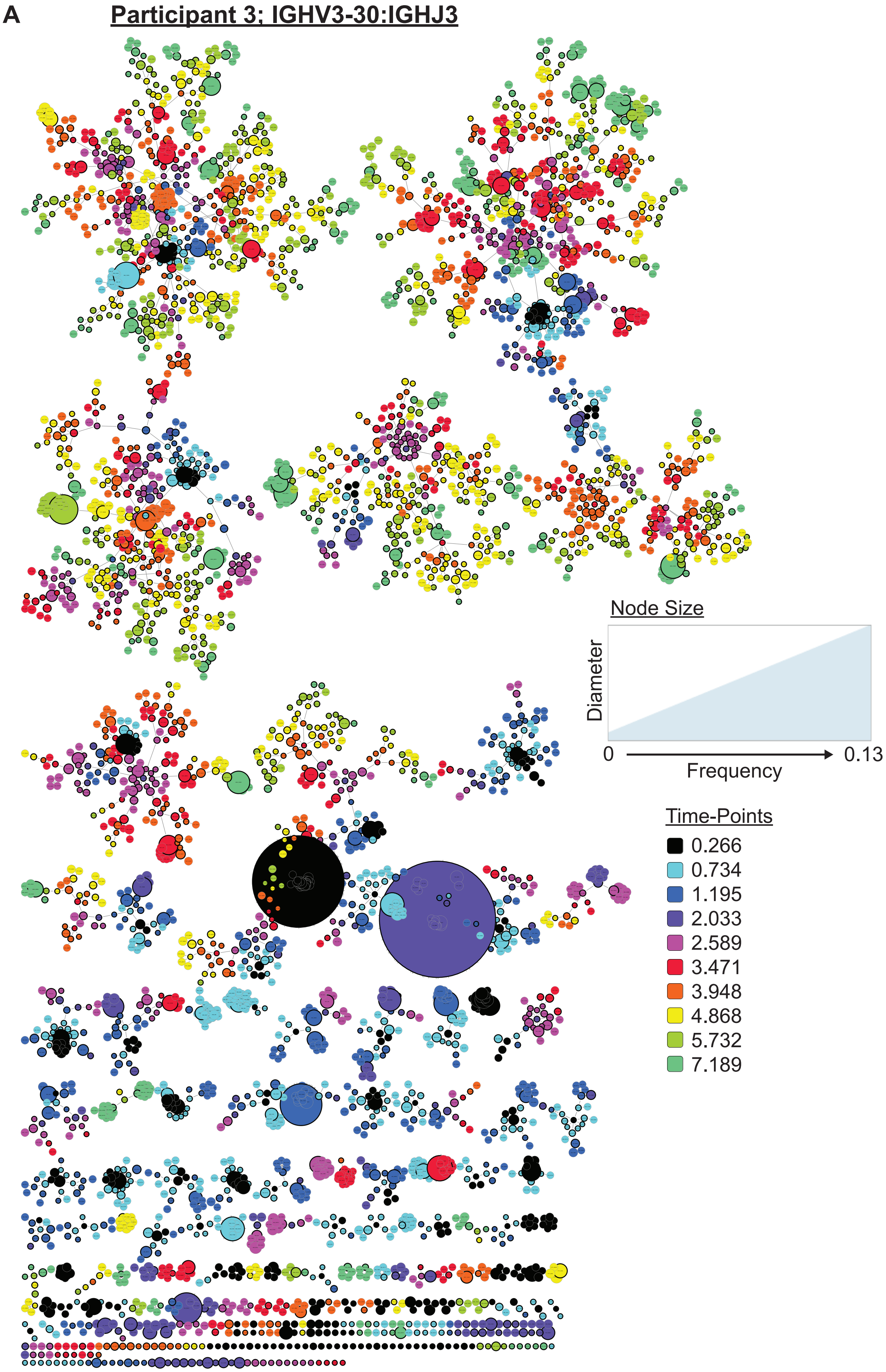
A
Participant 3; IGHV3-30:IGHJ3
Node Size
Diameter
0
Frequency
0.13
Time-Points
0.266
0.734
1.195
2.033
2.589
3.471
3.948
4.868
5.732
7.189

**B** **Participant 7; IGHV4-31:IGHJ5**

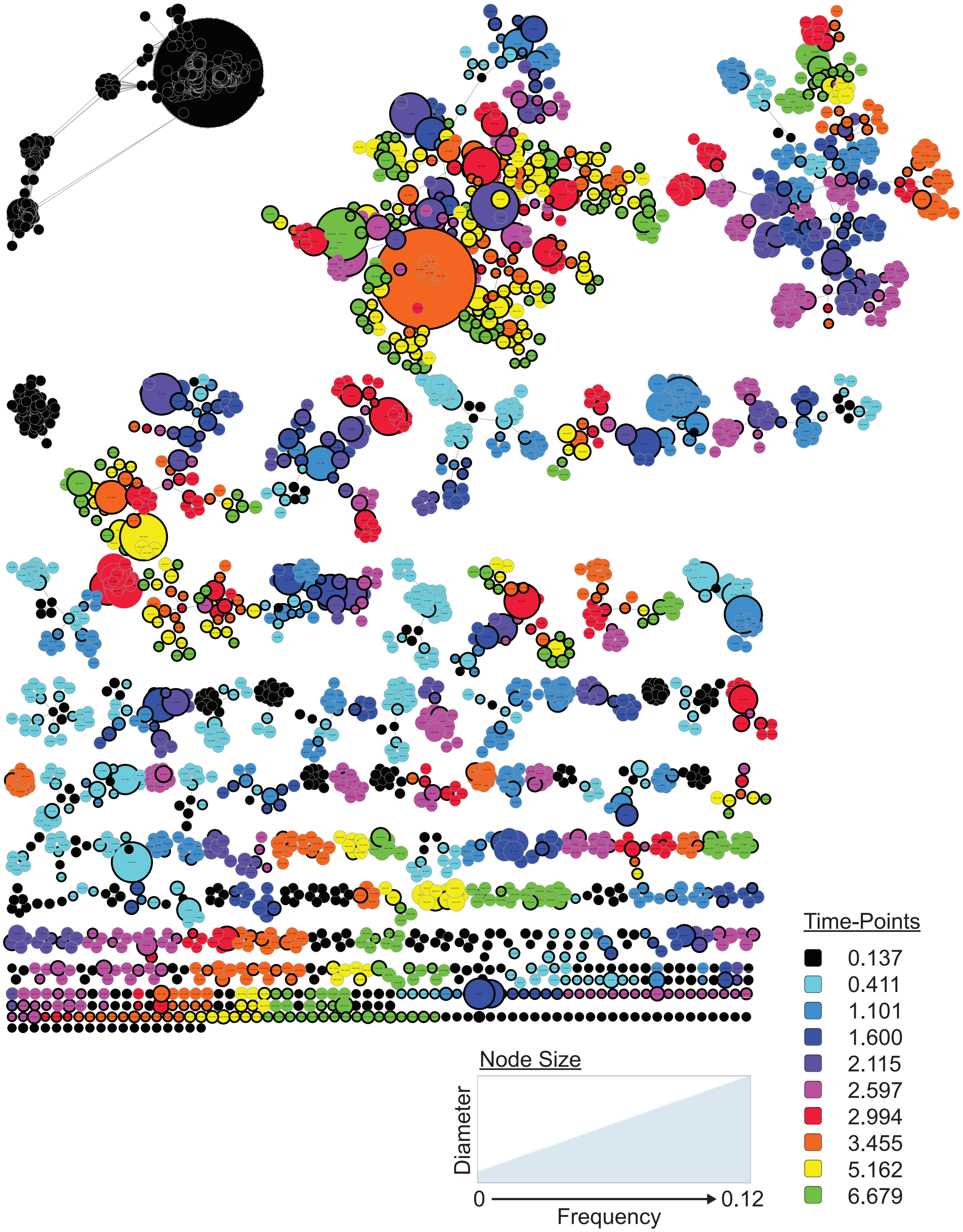

C

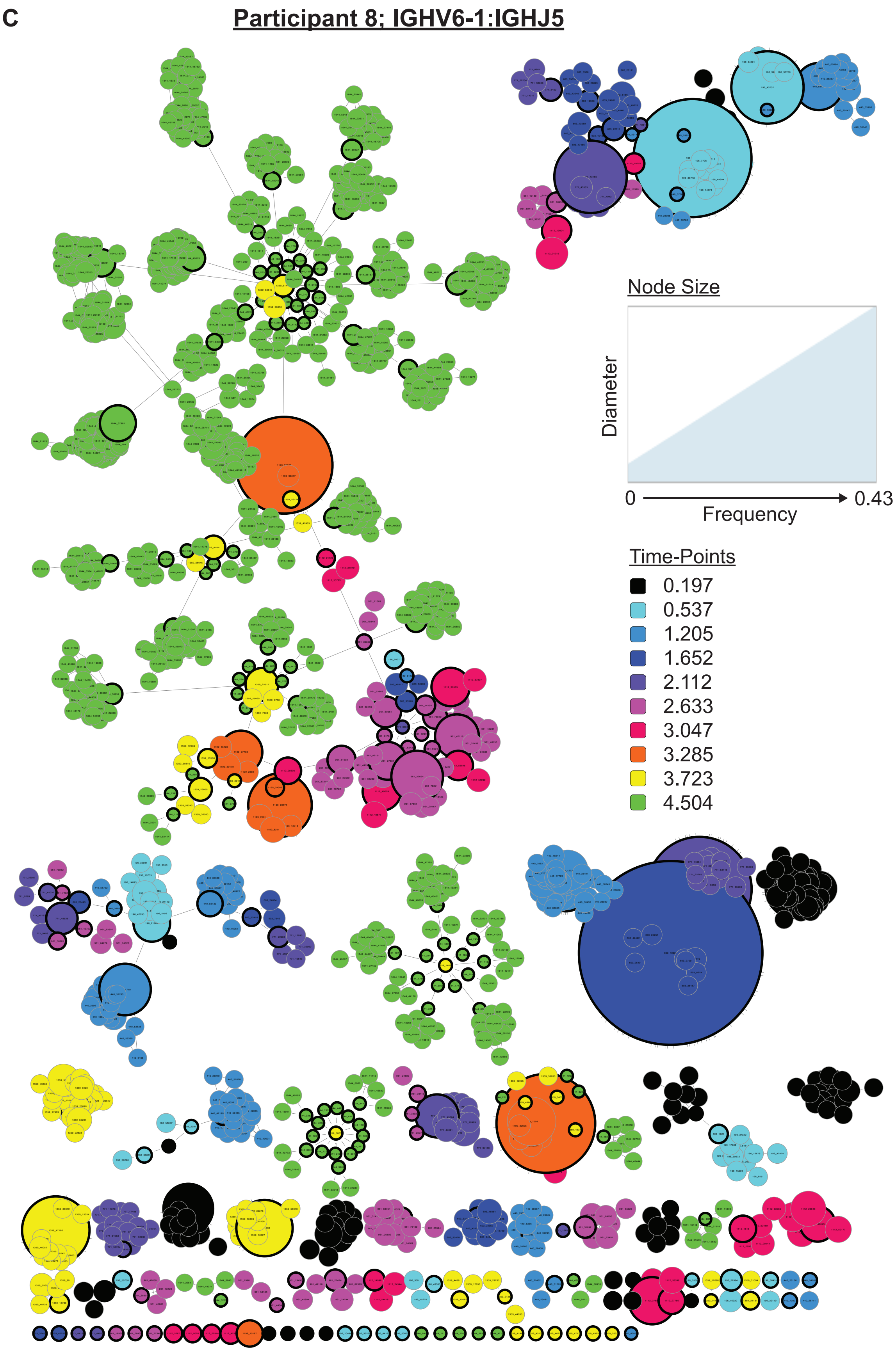

Participant 8; IGHV6-1:IGHJ5
Node Size
Diameter
0
Frequency
0.43
Time-Points
0.197
0.537
1.205
1.652
2.112
2.633
3.047
3.285
3.723
4.504

**Figure S16. Network plots of AbR lineages.**
Depicts the structure of each Ab lineage within a select HIV-associated AbR partition. Each node represents a unique sequence, and the size of a given node reflects the frequency of that sequence (within the AbR partition), at a given time-point. The fill color of a node reflects time-point, and if a node has a thicker black border then it was assigned to be a representative sequence of a sequence-cluster (borders are otherwise thinner and gray). Nodes of the same color (i.e. sequences in the same time-point) are linked with an edge if they were in the same cluster. Nodes that have different colors are linked with an edge if they were assigned to the same lineage. This inter-time-point linking only occurs between representative sequences. Taken together, each isolated grouping of nodes shows a family of related lineages. (A) AbR partition IGHV3-30:IGHJ3 in participant 3. (B) AbR partition IGHV4-31:IGHJ5 in participant 7. (C) AbR partition IGHV6-1:IGHJ5 in participant 8.

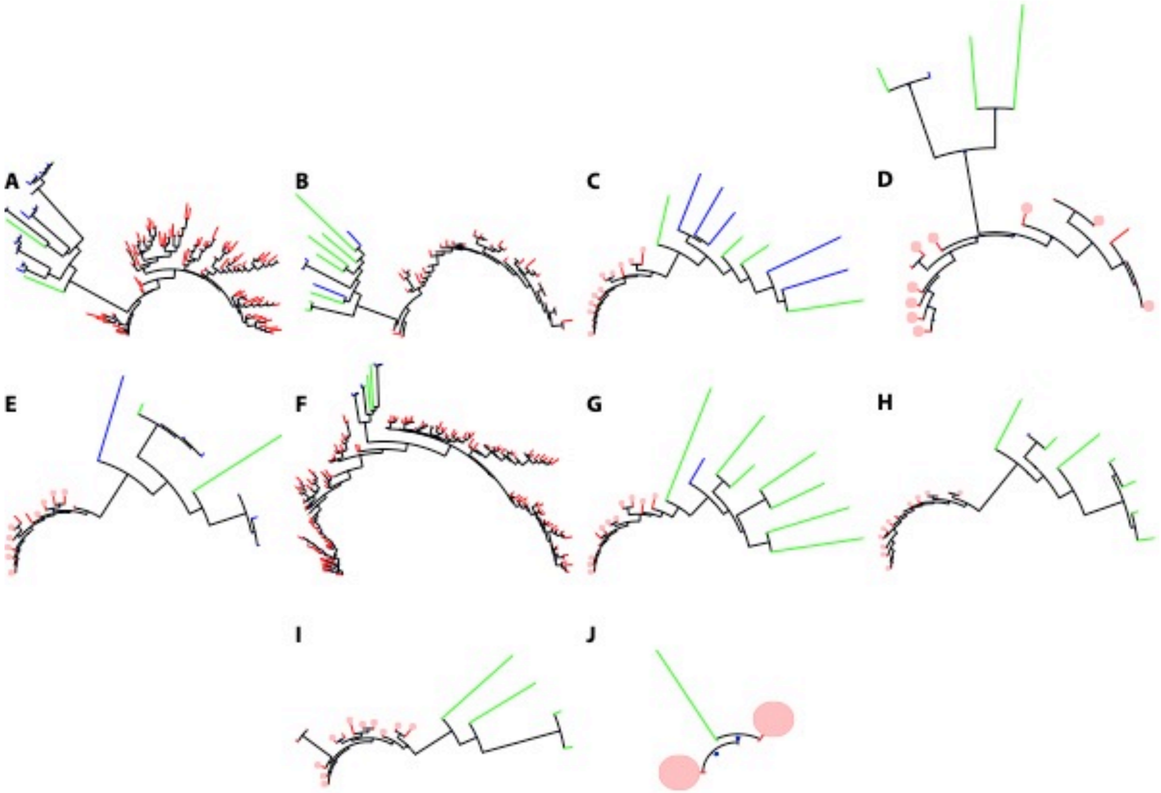


**Figure S17. Outlier HIV sequences.**
Phylogenetic trees depicting the relationship of HIV sequences for each participant. Each leaf on a tree represents a cluster of sequences, and the size of the circle at the terminus of a leaf gives the relative frequency of that cluster at a given time-point. Leaves that are colored blue were identified as outliers because they were found to have a closer genetic distance to clusters from another participant then clusters within their assigned participant (i.e. cross-contaminants). Leaves that are green were identified as outliers because they group closest with leaves identified a cross-contaminants.

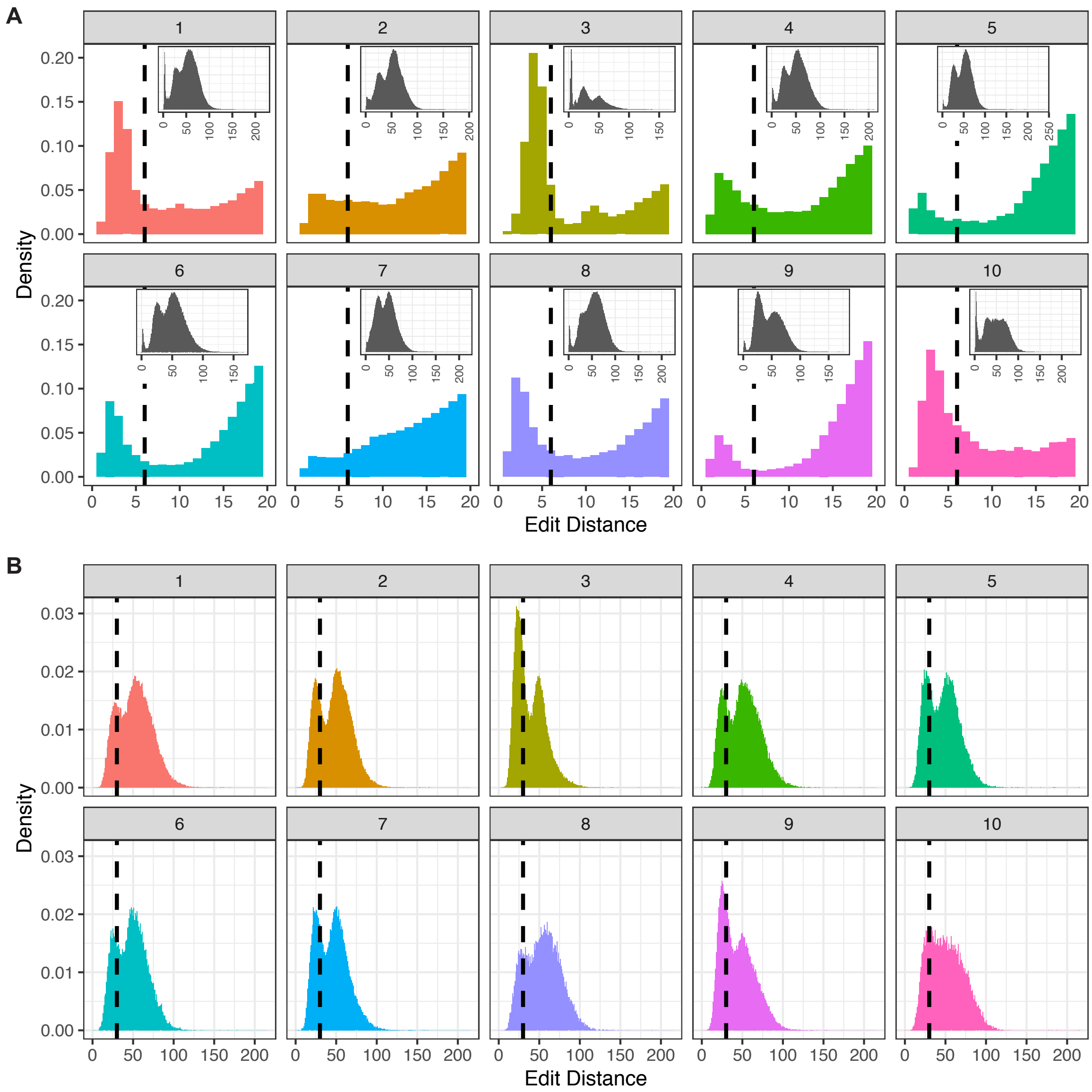


**Figure S18. Genetic distance distributions for AbR sequences.** Histograms of edit distance values between pairs of antibody sequences in the AbR. Top label foreach panel and color correspond to participant. (A) Histograms of edit distance values for all sequences within a time-point. The within time-point distance values are pooled across all time-points to generate the shown histograms. The main plots give a zoomed-in view for edit distance values 0-20. Insets show the entire histogram. Vertical dashed lines at x=6 show the edit distance cutoff that was used to assign sequences to the same cluster. (B) Histograms of edit distance values between adjacent time-points. This shows the edit distance distributions between pairs of representative cluster sequences across adjacent time-points. Vertical dashed lines at x=30 show the cutoff that was used to assign sequence clusters as being members of the same lineage.

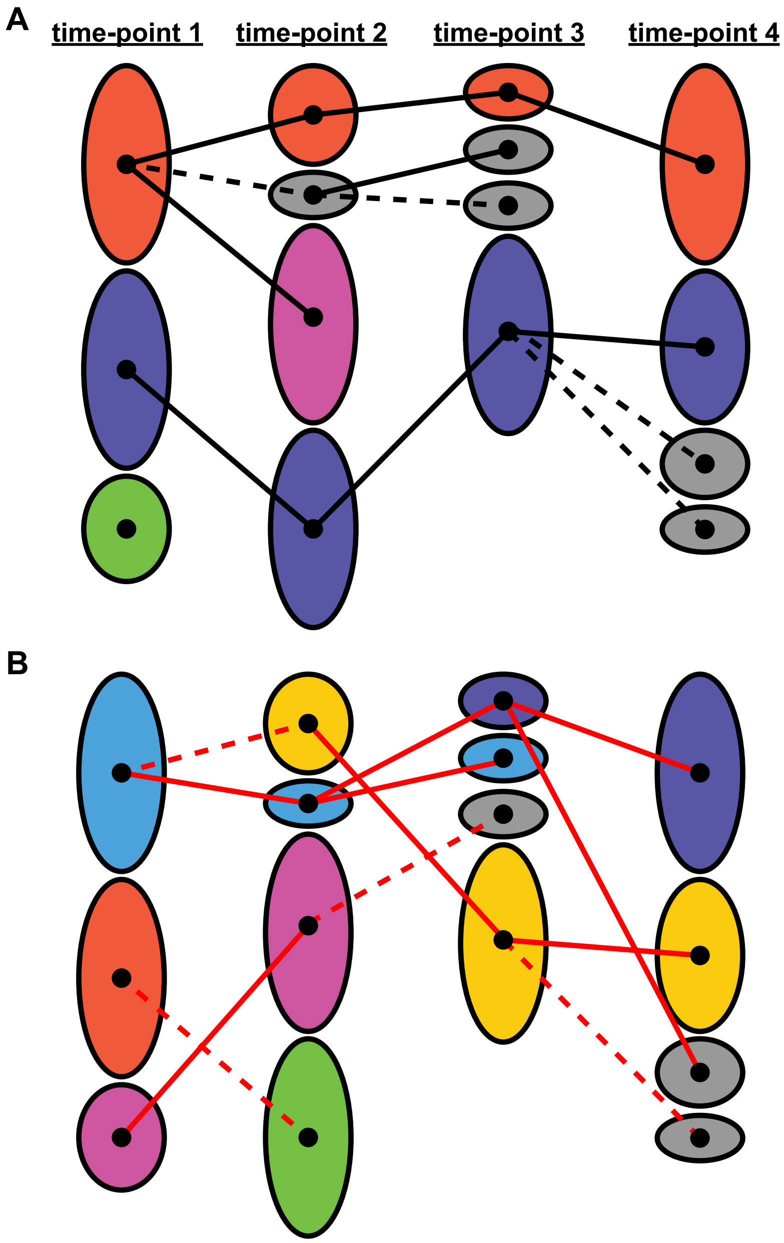


**Figure S19. Schematic of lineage clustering algorithm.**
Each ellipse represents a cluster of sequences, and the dot in the middle is the representative sequence for a given cluster. Dashed lines, or edges, represent lineage connections that fall above the edit distance threshold, while solid edges fall below. Clusters are colored by which lineage they belong to. In the cases of multifurcations, the parental color is assigned to the daughter cluster that has the lowest edit distance to the parent cluster. However, this is only pertinent for visualizations, such as this figure and Muller plots. Grey denotes lineages that do not meet the minimum frequency threshold. (A) Schematic of an observed lineage structure. The black edges linking clusters across time-points are formed by finding the cluster in the previous time-point that has the representative sequence with the smallest genetic distance to a given query representative

sequence. (B) Schematic of a simulated lineage structure. Red edges linking clusters across time-points are determined randomly.

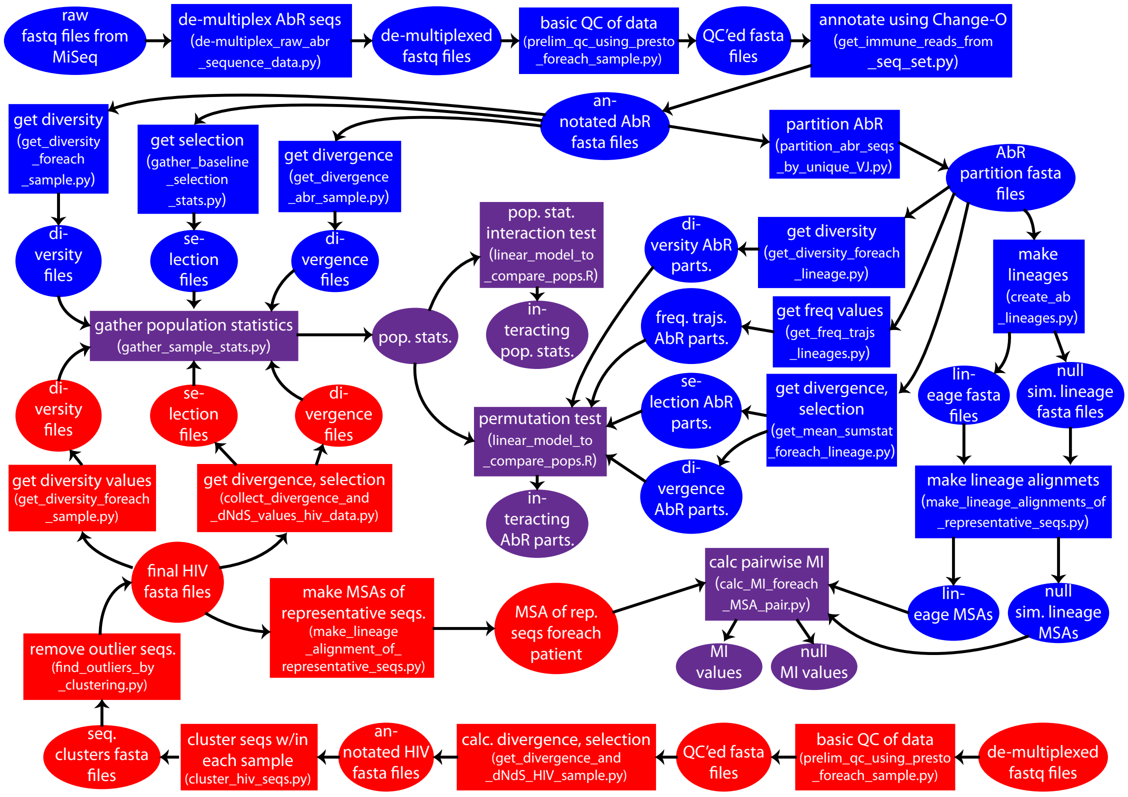


**Figure S20. Flow chart for entire project.**
Each node represents a step in the project, where ellipses represent data files (ex. fastq, fasta, txt, etc.), and rectangles represent data processing events (i.e. scripts). Blue nodes show steps that only have/use AbR data, red nodes show steps that only have/use HIV data, and purple nodes show steps where AbR and HIV data have been merged or compared. The AbR pipeline begins at the top-left of the chart, and the HIV pipeline begins at the bottom-right. The text between parentheses in each rectangular node gives the name of the script that was used to complete the corresponding data processing event. This can be used a reference should one want to use our code.

### Supplementary Tables

| Gene Name | V Gene Score | HIV Literature Dataset Count | Flu Literature Dataset Count |
|---|---|---|---|
| IGHV4-59 | 5.692163182 | 5 | 71 |
| IGHV1-18 | 5.135748975 | 1 | 14 |
| IGHV3-23 | 4.069059968 | 0 | 48 |
| IGHV3-30-3 | 3.996480894 | 0 | 0 |
| IGHV4-39 | 3.931435639 | 7 | 37 |
| IGHV3-30 | 3.826571542 | 14 | 18 |
| IGHV4-61 | 3.825996333 | 0 | 1 |
| IGHV1-46 | 3.708051547 | 0 | 0 |
| IGHV1-8 | 3.705576595 | 18 | 0 |
| IGHV3-21 | 3.301116439 | 1 | 40 |
| IGHV3-11 | 3.233458041 | 0 | 2 |
| IGHV3-33 | 3.020795329 | 1 | 3 |
| IGHV3-66 | 2.99037641 | 0 | 0 |
| IGHV3-74 | 2.801852721 | 0 | 0 |
| IGHV3-7 | 2.769498585 | 0 | 19 |
| IGHV4-34 | 2.036779778 | 3 | 7 |
| IGHV1-2 | 2.029337852 | 18 | 7 |
| IGHV1-3 | 2.018780447 | 1 | 0 |
| IGHV6-1 | 1.922879261 | 0 | 0 |
| IGHV1-69 | 1.809503004 | 2 | 108 |
| IGHV2-5 | 1.723749802 | 1 | 8 |
| IGHV4-4 | 1.672966404 | 0 | 10 |
| IGHV3-15 | 1.532535707 | 1 | 2 |
| IGHV3-9 | 1.532147253 | 0 | 5 |
| IGHV3-53 | 1.446589506 | 0 | 1 |
| IGHV1-24 | 1.298021556 | 0 | 0 |
| IGHV7-4-1 | 1.217541317 | 0 | 2 |
| IGHV5-51 | 1.212203964 | 0 | 7 |
| IGHV3-48 | 1.206518076 | 0 | 9 |
| IGHV4-30-2 | 1.189591763 | 0 | 0 |
| IGHV3-13 | 1.123988038 | 0 | 6 |
| IGHV3-64 | 0.851467542 | 0 | 1 |
| IGHV4-30-4 | 0.790029158 | 0 | 3 |
| IGHV4-31 | 0.708777704 | 0 | 2 |
| IGHV3-72 | 0.298207811 | 0 | 0 |
| IGHV2-70D | 0.220188607 | 0 | 0 |
| IGHV3-43 | 0.209900779 | 0 | 0 |
| IGHV2-70 | 0.115635873 | 0 | 1 |
| IGHV4-38-2 | 0.103112693 | 0 | 0 |
| **Total** | NA | **73** | **432** |

**Table S1. Literature datasets.**
